# Modulation of Aβ_1-42_ Aggregation by a SARS-COV-2 Protein Fragment

**DOI:** 10.1101/2025.08.08.669385

**Authors:** Malinda B. Premathilaka, Ulrich H. E. Hansmann

**Affiliations:** Department of Chemistry & Biochemistry, University of Oklahoma, Norman, OK 73019, USA

**Author notes:** **Corresponding Author: Ulrich H.E. Hansmann** - Department of Chemistry & Biochemistry, University of Oklahoma, Norman, Oklahoma 73019, United States.

**Keywords:** Molecular Dynamics Simulations, SARS-COV-2, Aβ, Amyloids

## Abstract

A number of studies have pointed out to the possibility that SARS-COV-2 infections could trigger amyloid diseases such as Parkinson’s disease or type-II diabetes. In the present study we probe this question for Alzheimer’s disease which is connected with presence of amyloids rich in Aβ-peptides. For this purpose, we study by way of molecular dynamics simulations the interaction between the fragment FKNIDGYFKI of the Spike protein with Aβ_1-42_ monomer and two fibril models, one patient-derived and one synthetic. Our results are compared with previous studies of other amyloid-forming proteins to identify commonalities and differences in the modulation of amyloid-formation by the viral protein fragment.

**Table of Contents Figure:** 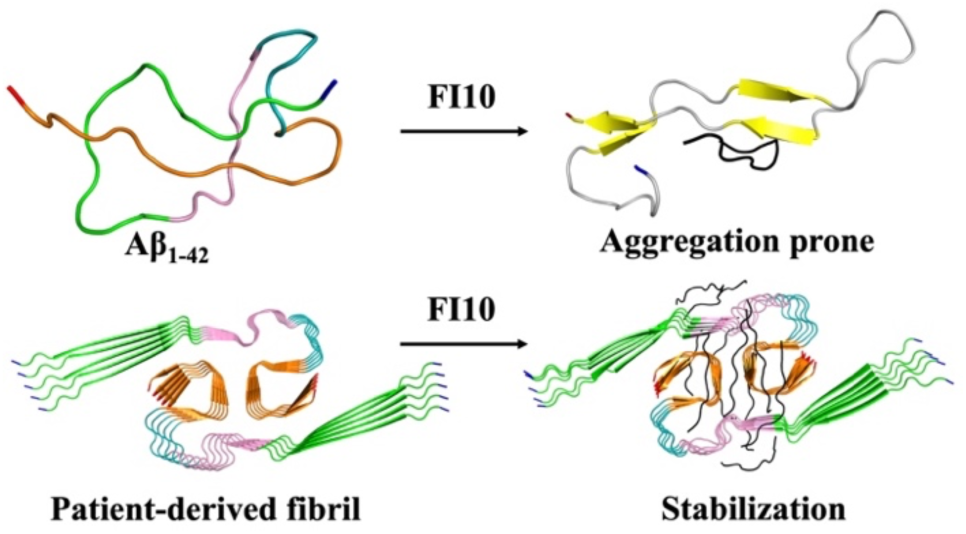

## Introduction

Presence of amyloids in humans is often associated with neurodegenerative and other diseases^1^, but there is only an incomplete understanding of how and by what agents amyloids form and propagate into fibrils. Various studies have shown that *in vitro* microbial proteins can seed aggregation of Aβ and α-synuclein (αS).^2–9^ For instance, SARS-COV-2-triggered amyloid formation is seen *in vitro* for αS^6^, implicated in Parkinson Disease (PD); and correlations between COVID-19 and outbreak of PD have been observed.^10,11^ Using extensive molecular dynamic simulations, we could show that amyloidogenic SARS-COV-2 protein fragments can enhance aggregation of αS by shifting the ensemble of monomers toward more aggregation-prone conformations and by differentially stabilizing toxic fibril polymorphs.^12,13^ As such short fragments are cleaved from SARS-COV-2 proteins by enzymes released from neutrophils during acute inflammation, and form *in vitro* amyloids,^14^ our results therefore point to a potential mechanism by which COVID-19 may cause PD.

In terms of the number of patients, a much more devasting neurodegenerative disease is Alzheimer’s disease (AD) connected to presence of amyloids rich in Aβ peptides. As correlations between COVID-19 and AD appear to exist^15^, it is possible that the presence of SARS-COV-2 in the brain^16–18^ ultimately raises the risk for AD. This conjecture is supported by the observation that glycoproteins on herpes simplex virus 1 (HSV-1) catalyze aggregation of Aβ.^3,19^ Similarly, the HIV-TAT protein induces Aβ amyloid formation and propagation into neurotoxic fibrils by forming complexes with Aβ.^20^ Hence, it is important to study whether and how viral proteins from SARS-COV-2 can also seed or modulate aggregates of Aβ proteins.

This is the purpose of the present study where we investigate the effect of the SARS-COV-2 specific FI10 peptide, released after cleavage by the enzyme neutrophil elastase from the spike protein (the segment 194-203: FKNIDGYFKI), on the ensemble of conformations of Aβ_1-42_ monomers. This task is eased by the wealth of computational and experimental data for Aβ_1-42_ monomers and fibrils that can help us to evaluate our results. Our goal is to probe whether the monomer ensemble is shifted towards conformations that have a higher propensity for aggregation. The end product of the Aβ aggregation process is a polymorphic set of fibrils, whose structures do not only depend on experimental conditions but are often also correlated with severity and disease propagation of AD. For this reason, we also consider the effect of FI10 on the stability of two experimentally resolved fibril models: a patient-derived one with Protein Data Bank (PDB) identifier 7Q4B and a synthetic fibril model that has PDB-ID 5KK3. Besides the potential health relevance of documenting an aggregation-modulating effect of SARS-COV-2 proteins on Aβ_1-42_ peptides, we try to establish a general mechanism for pathogen-induced amyloid formation by comparing the results for Aβ with our earlier work studying the effect of SARS-COV-2 protein fragments on the aggregation of αS, SAA and amylin.^12,13,21,22^

## Materials and Methods

### System Preparation

We rely on molecular dynamics (MD) simulations to study how the ten-residue segment FKNIDGYFKI (FI10) of the Spike protein from SARS-COV-2 affects the conformational ensemble of Aβ_1-42_ monomer, and whether the observed changes suggest an increased propensity for aggregation. While Aβ_1-40_ peptides are more frequent, we chose for this investigation Aβ_1-42_ peptides as they are associated with more severe forms of AD. The initial configurations for the Aβ_42_ monomer were obtained from one of our previous works, which employed our Resolution Exchange with Tunneling (ResET) to sample Aβ_1-42_ monomer conformations. In order to be consistent with our earlier work^23^, we add again at the N- and C-terminal a acetyl and a methyl group.

Simulations starting from the Aβ_1-42_ monomer described above serve as control in our study. For investigating the effect of SARS-COV-2 protein fragments on Aβ_1-42_ we have performed simulations that start from configurations where a FI10 peptide bind to this model. Following Nyström and Hammarström^14^ we extracted the residues **F**194-**I**203 from the resolved oSARS-COV-2 Spike protein model (PDB ID: 6VXX) as the initial configuration of the FI10 viral fragment, with the N- and C-terminal of the peptide capped again by a NH_3_^+^ and COO^-^ group. Using HADDOCK^24, 25^ we docked the FI10 peptide in a ratio of 1:1 with the Aβ_1-42_ monomer. The resulting configurations are shown in **Supplemental Figure SF1**. Note that the FI10 peptide is not constraint to this initial position but can move freely and over the course of the simulation even detach from the Aβ_1-42_ monomer.

Besides probing the interaction with Aβ_1-42_ monomers, we also looked into the effect of the viral protein fragment FI10 on fibrils formed from Aβ_1-42_ chains. As there is a considerable polymorphism in the resolved structures deposited in the Protein Data Bank (PDB), we have considered in this study two distinct fibril models: the patient derived Aβ42 fibril model of type I (PDB ID: 7Q4B), which was resolved by cryo-EM for the segment of residues 9-42, and the synthetic fibril model (PDB ID: 5KK3) that has been derived *in vitro* and was resolved by NMR for residues 11-42. Comparing the two fibril models allows us to test if there is a differential modulation of fibril stability by the viral protein fragment. Both fibril structures have two-fold symmetry. The patient-derived fibril model is composed of five layers, while the synthetic fibril model is made of nine layers. To maintain the consistency among the fibril simulations, we therefore extracted the first five layers from the synthetic fibril structure for our model. As in both models the first N-terminal residues were missing reflecting disordered N-terminal regions, we added these residues with PyMol^26^, and capped the N and C terminals of the chains with acetyl and methyl groups, respectively. The so-obtained two-fold-five-layer fibril fragments were relaxed in 3 ns simulated annealing runs where the experimentally resolved parts of the fibril were restrained. For this purpose, the respective fibril fragment was heated up to 510 K within first 200 ps, then cooled down to 310 K within next 2600 ps, and finally equilibrated at 310 K over the 200 ps. The so obtained structures (shown in Figure 1e–1f) served as start configurations in our control simulations (i.e., in absence of FI10), and were compared with simulations where ten FI10 peptides are docked with the fibril fragments (shown also in **Supplemental Figure SF1**) in a ratio of 1:1 with the Aβ_1-42_ chains using again HADDOCK. ^24, 25^

**Figure 1.**
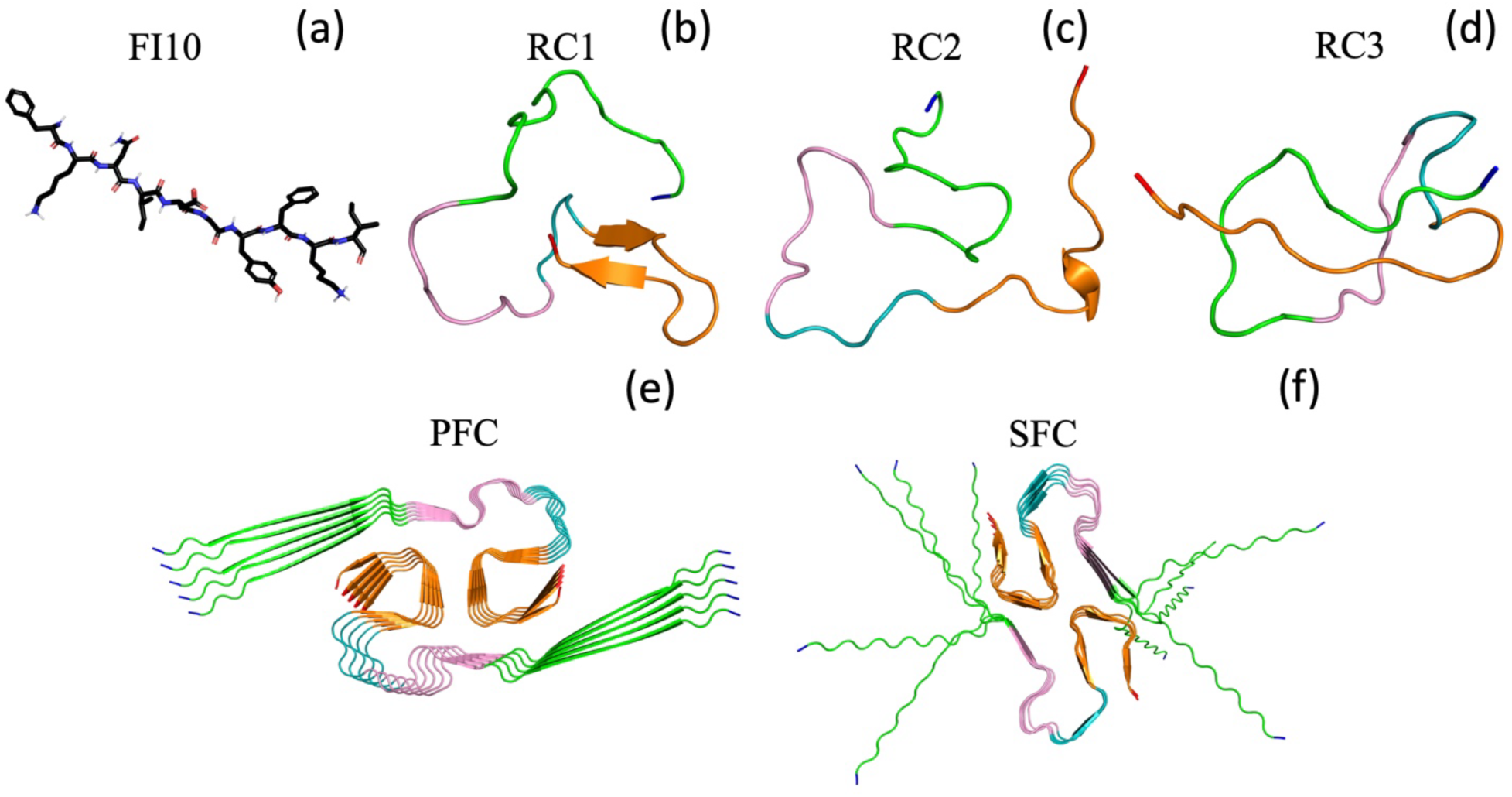
(a) Licorice representation of the SARS-COV-2 protein fragment FI10 used in this study. Ribbon representations of the three initial configurations of Aβ_1-42_ monomers (RC1, RC2, and RC3) are shown in (b-d), with the considered regions colored in green (region N, residues 1-15), pink (region M1, residues 16-24), teal (region M2, residues 25-29) and orange (region C, residues 30-42). The N- and C-terminals are colored in blue and red, respectively. Finally, we show sketches of the two decamer Aβ_1-42_ fibril models used in our study, the patient-derived model (PFC) with PDB-ID 7Q4B (e) and the synthetic fibril model (SFC) with PDB-ID 5KK3 (f).

### General Simulation Protocol

The set-up and simulation of all systems relies on the GROMACS 2022 package^27^. We use the CHARMM 36m all-atom force-field^28^ with TIP3P explicit water^29^ as implemented in the GROMACS package to describe the inter- and intramolecular interactions for the monomer and fibrils. Previous work performed in our group^30,31^ and in literature^32–34^ has shown that this force-field and water model combination is well-suited for simulations of fibrils and oligomers. Hydrogen atoms are added with the *pdb2gmx* module of the GROMACS suite^27^. The respective start configurations are put in the center of the cubic box, with at least 15 Å between the solute and the edge of the box. Periodic boundary conditions are employed. The systems are solvated with water molecules, and counterions are added to neutralize the system, with the Na^+^ and Cl^-^ ions at a physiological ion concentration of 150 mM NaCl. Both the number of water molecules and the total number of atoms are listed in **Table 1**. The energy of each system is minimized by steepest decent for up to 50,000 steps, and afterwards the system is equilibrated at 310 K for 200 ps at constant volume and in an additional 200 ps at constant pressure (1 bar), constraining the positions of heavy atoms with a force constant of 1000 kJ mol^-1^ nm^-2^.

**Table 1.**
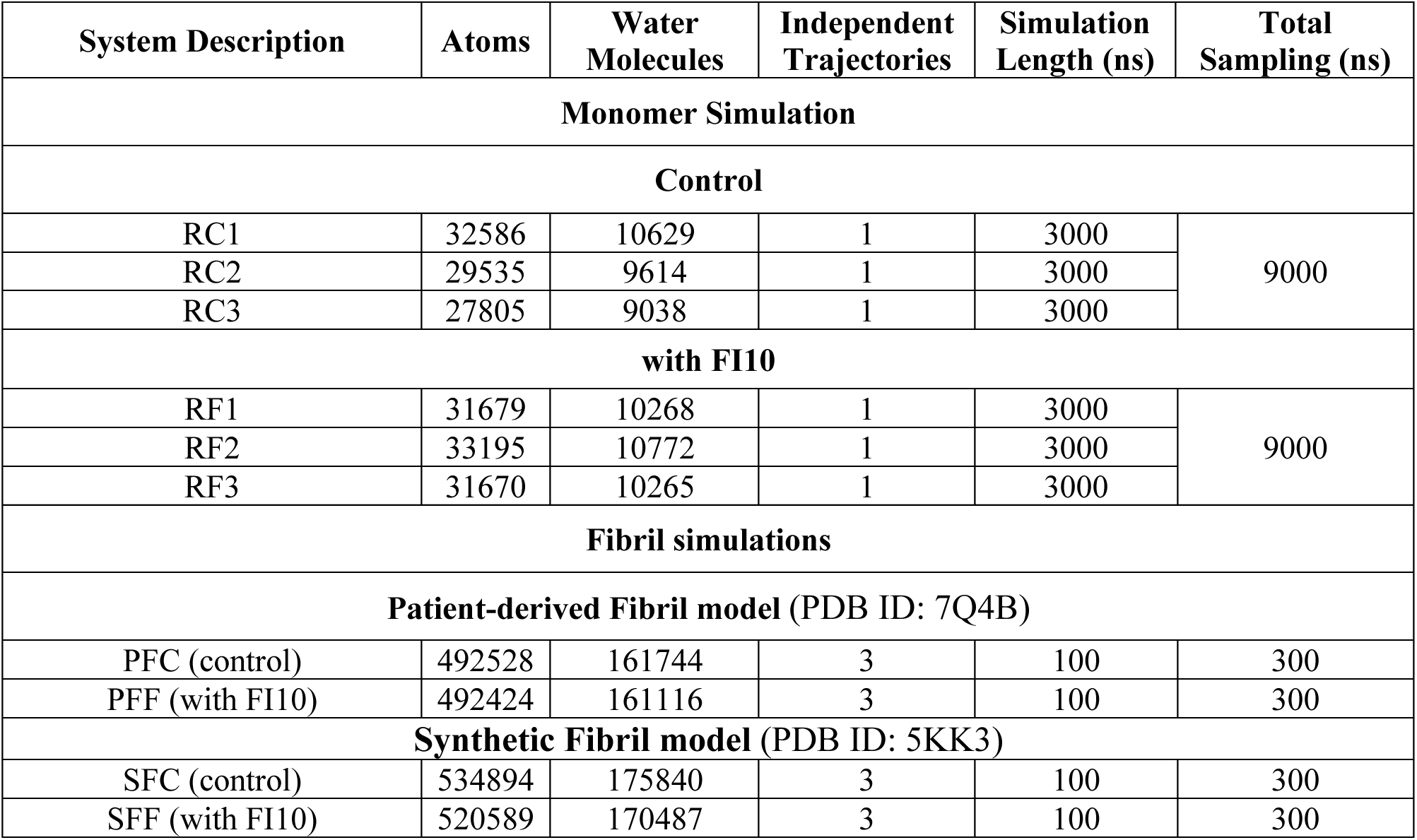
Simulated systems.

After the equilibration, our monomer simulations are followed at constant temperature (310 K) and constant pressure (1 bar) over three µs for each replica. The fibril simulations run at the same temperature and pressure, but because of the much larger system size only for 100 ns. We used v-rescale thermostat^35^ with 0.1 ps coupling constant to control the temperature. The Parinello-Rahman barostat^36^ with 2 ps relaxation time was employed for the pressure coupling. The SETTLE algorithm^37^ was used to maintain the rigidity of the molecules, while LINCS algorithm^38^ was applied to restrain the proteins bonds and kept hydrogen atoms to their equilibrium distance. In our simulations, we used 2 fs time step to integrate the equation of motion. We employed particle-mesh Ewald (PME) method^39^ with 12 Å real space cutoff and 1.6 Å Fourier grid spacing to deal with long-range electrostatic interactions. A cutoff of 12 Å was introduced to deal with short-range interactions. **Table 1** shows the lengths and total sampling durations in our simulations. Atomic coordinates of the initial and final configurations of all trajectories are provided as **supplemental material**.

### Trajectory Analysis

We used VMD^40^, PyMol^26^, and UCSF Chimera^41^ to visualize our trajectories and conformations. GROMACS tools were employed to calculate the root mean square deviation (RMSD), root mean square fluctuation (RMSF), solvent accessible surface area (SASA), and radius of gyration (Rg). The solvent accessible surface area was calculated using a 1.4 Å spherical probe. We performed intrastrand contact analysis using MDTraj library^42^ and 4.5 Å cutoff between the closest heavy atoms in a residue pair. The secondary structures of the peptides were determined using MDAnalysis library^43^ and the Dictionary of Secondary Structure in Proteins (DSSP) algorithm.^44^ Salt bridges were analyzed using MDAnalysis with a cutoff of 4.0 Å.

We performed a cluster analysis for the monomer simulations to identify the most representative conformations during the simulation using the method explained by Daura et al.^45^ with a cutoff of 0.5 nm for the backbone RMSD calculation in cluster analysis. The number of contacts between FI10 and Aβ_1-42_ was calculated using the MDAnalysis library. We used the same cutoff (4.5 Å) as for calculating the intra-strand contacts in order to calculate the contacts between Aβ_1-42_ and FI10.

## Results and Discussion

### Monomer simulations

Aβ_1-42_ is an intrinsically disordered peptide. While models of Aβ_1-42_ peptides with partially stable secondary structures have been deposited in the Protein Data Bank (for instance, the helix-rich model with PDB-ID 1Z0Q), these models reflect experimental conditions that stabilize secondary structure, and are not necessarily representative for the peptide in water. For this reason, we use as start configurations for our control simulations (without FI10 bound to it) equilibrium configurations taken after 1 μs from trajectories obtained in previous all-atom molecular dynamics simulations^23^ (see the Methods section). As expected for such equilibrium configurations, all three are disordered with little secondary structure (in R1 β-strands for segments I29-A31 and V39-I41, in R2 a short helical segment L34-V36, and only coil structure in R3).

Over the course of the trajectories, quantities such as the radius of gyration, solvent accessible surface area or number of intra-strand contacts fluctuate around an equilibrium value, see **Figure 2a) - 2c)**. However, it is not obvious that these three initial configurations represent also equilibrium structures for the complex formed by Aβ_1-42_ with FI10, and some time may be needed to approach equilibrium. For this reason, we chose for our analysis only the last 2μs of our trajectories at which time the root-mean-square-deviations to the start (not shown) had reached a plateau for all systems. As observed similarly in previous work for αS, presence of the viral protein fragment FI10 shifts the ensemble of Aβ_1-42_ monomer configurations to more stretched and potentially more aggregation-prone ones with larger radius of gyration (Rg) and solvent accessible surface area (SASA), but lower number of intra-strand contacts, see **Figure 2d) - 2f)**.

**Figure 2.**
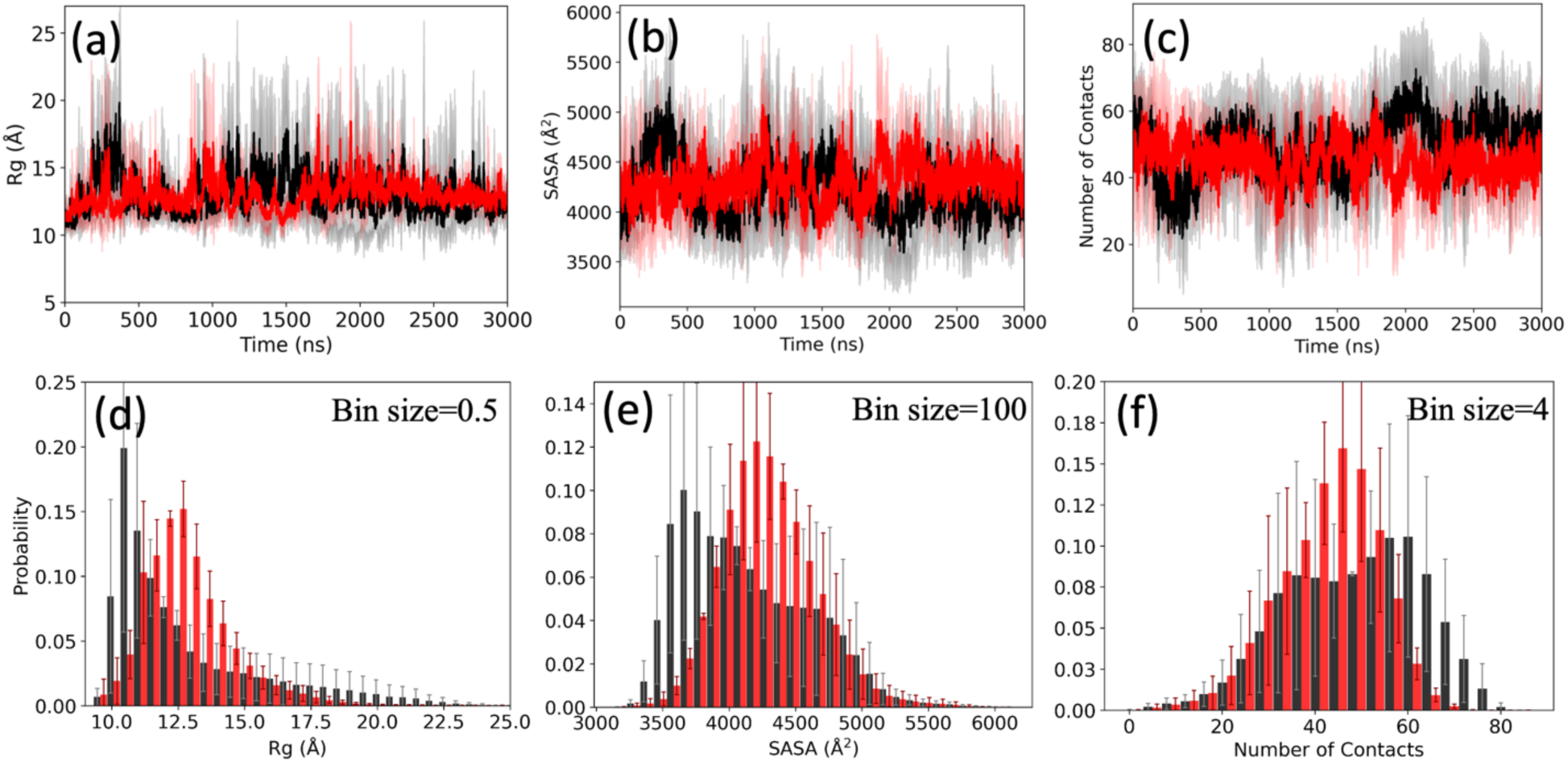
The (a) radius of gyration (Rg), (b) solvent accessible surface area (SASA), and (c) the number of contacts (*n*_C_) of the Aβ_1-42_ monomer in the absence (black) and presence (red) of FI10 as a function of time. Shown is the difference to the corresponding value at time *t=0*. The plotted values are averages over three independent trajectories and the shaded regions mark the standard deviation of the averages. In (d), (e) and (f) we show the corresponding normalized averaged distributions of the sampled data collected over the final 2.0 μs of each trajectory.

These differences between simulations in absence and presence of FI10 seem to be associated with a lower flexibility of the C-terminal residues 30-42 leading to increased values for the residue-wise root mean-square fluctuations shown in **Figure 3a** that are only partially correlated with an increased binding propensity of FI10 to C-terminal Aβ_1-42_ residues in **Figure 3b**. On the other hand, the time series of residue-wise secondary structure for all six trajectories in **Figure 4** seems to indicate a higher strandness in the C-terminus when interacting with FI10 (**Figure 4b**) over what is seen in the control (**Figure 4a**), however, this signal is weak.

**Figure 3.**
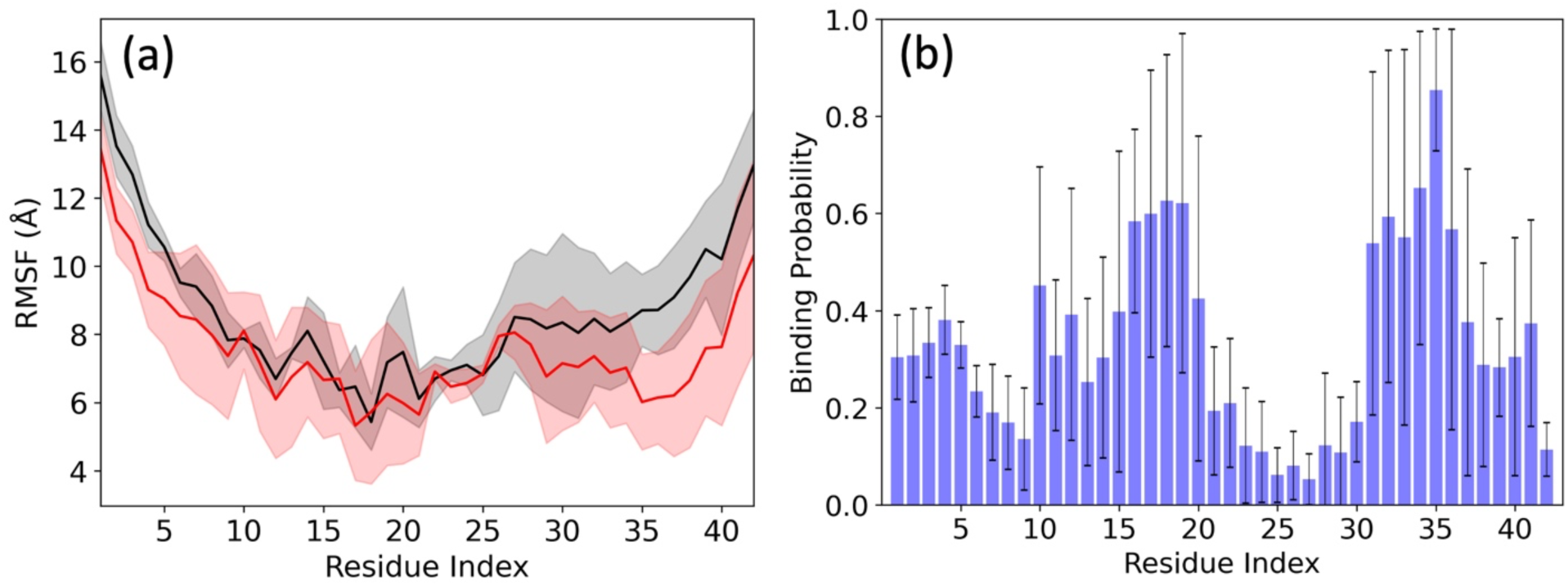
(a) Residue-wise mean square fluctuation (RMSF; in Å) obtained from Aβ_1-42_ monomer simulations in the absence (black) and presence (red) of the FI10 segment. binding probability of the FI10 segment to the Aβ_1-42_ monomer residues is shown in (b). Data are averaged over the final two μs of each trajectory. Shaded regions in (a) mark the standard deviation of the averages.

**Figure 4.**
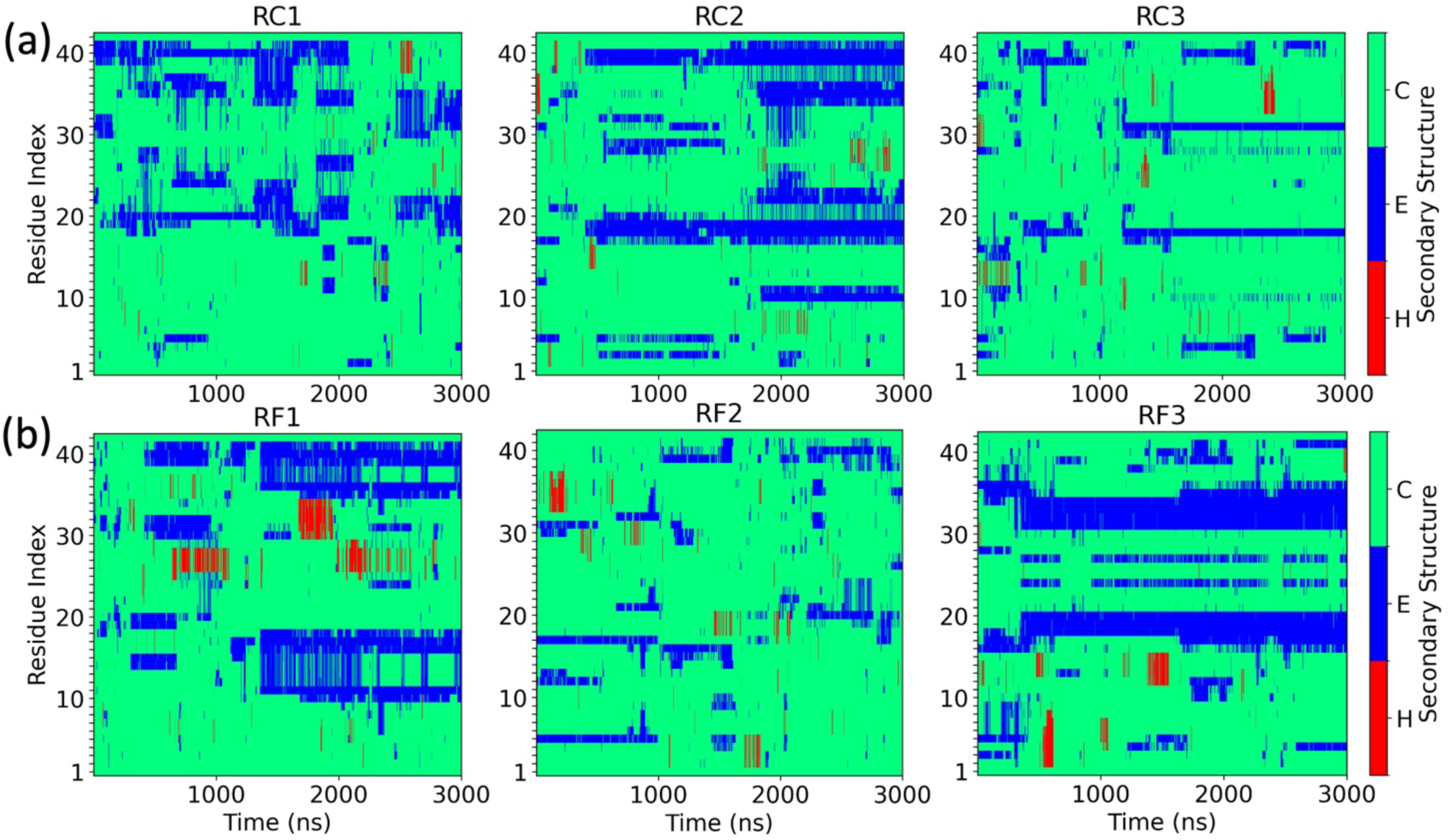
Time series of the residue-wise secondary structure (helix (H), strand (E) and coil (C)) for (a) the three trajectories where FI10 is absent, and (b) the three trajectories where FI10 can interact with the Aβ_1-42_ monomer.

In order to get a deeper understanding of the effect of FI10 on the ensemble of Aβ_1-42_ monomers, we have clustered the configurations collected over the last 2 μs in the six trajectories (three for the control, and three with FI10 present) according to the procedure described in the method section. Considering only clusters containing more than 5% of the configurations in a trajectory, we are left in the control simulations with two clusters in trajectory 1, four in trajectory 2 and two in trajectory 3. The frequencies of these clusters are listed in **Table 2**. Centroids of these eight clusters, compromising together around 53% of the 120000 snapshots in all three trajectories, are shown in **Figure 5**.

**Figure 5.**
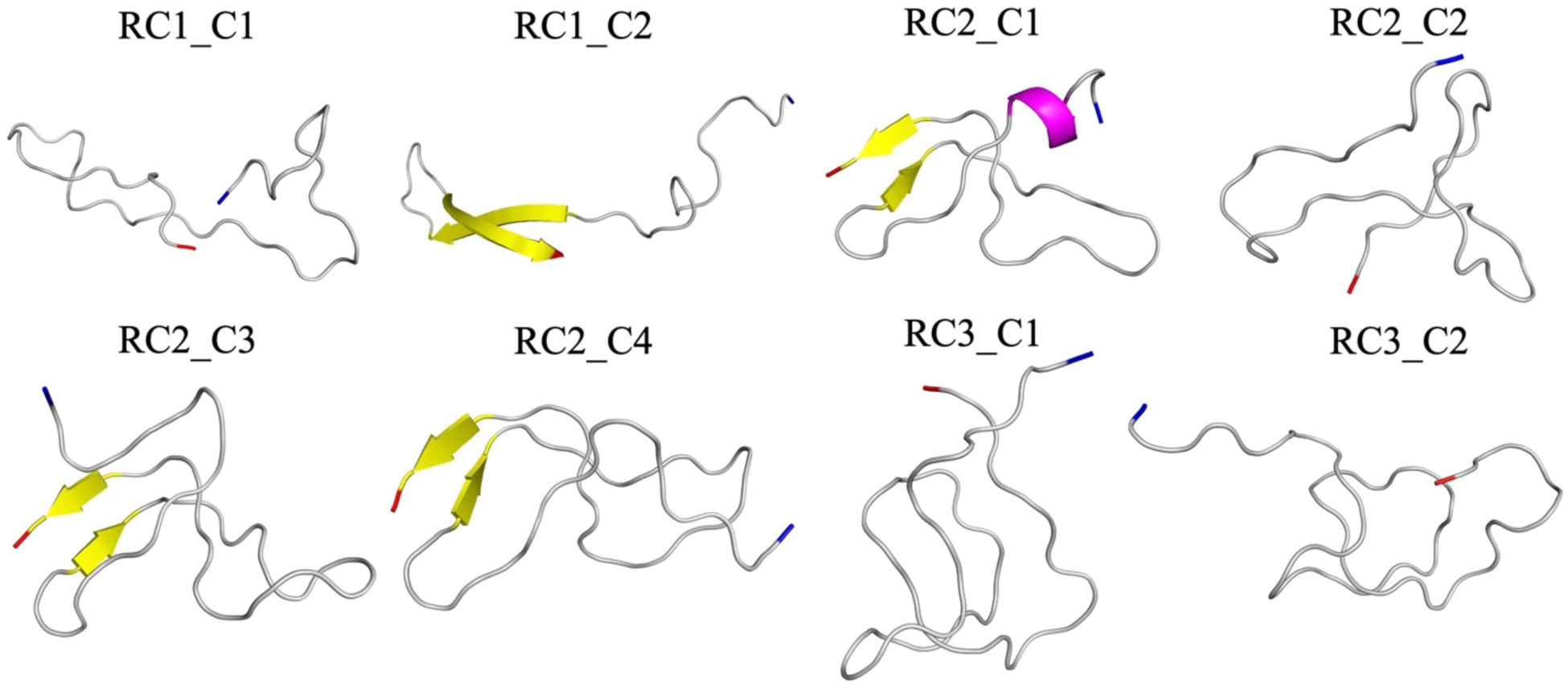
Ribbon representations of the centroid conformations in the eight most populated clusters of configurations sampled over the final 2 μs in the three control trajectories (where FI10 is absent). The N- and C-terminals are colored in blue and red, respectively.

**Table 2.**
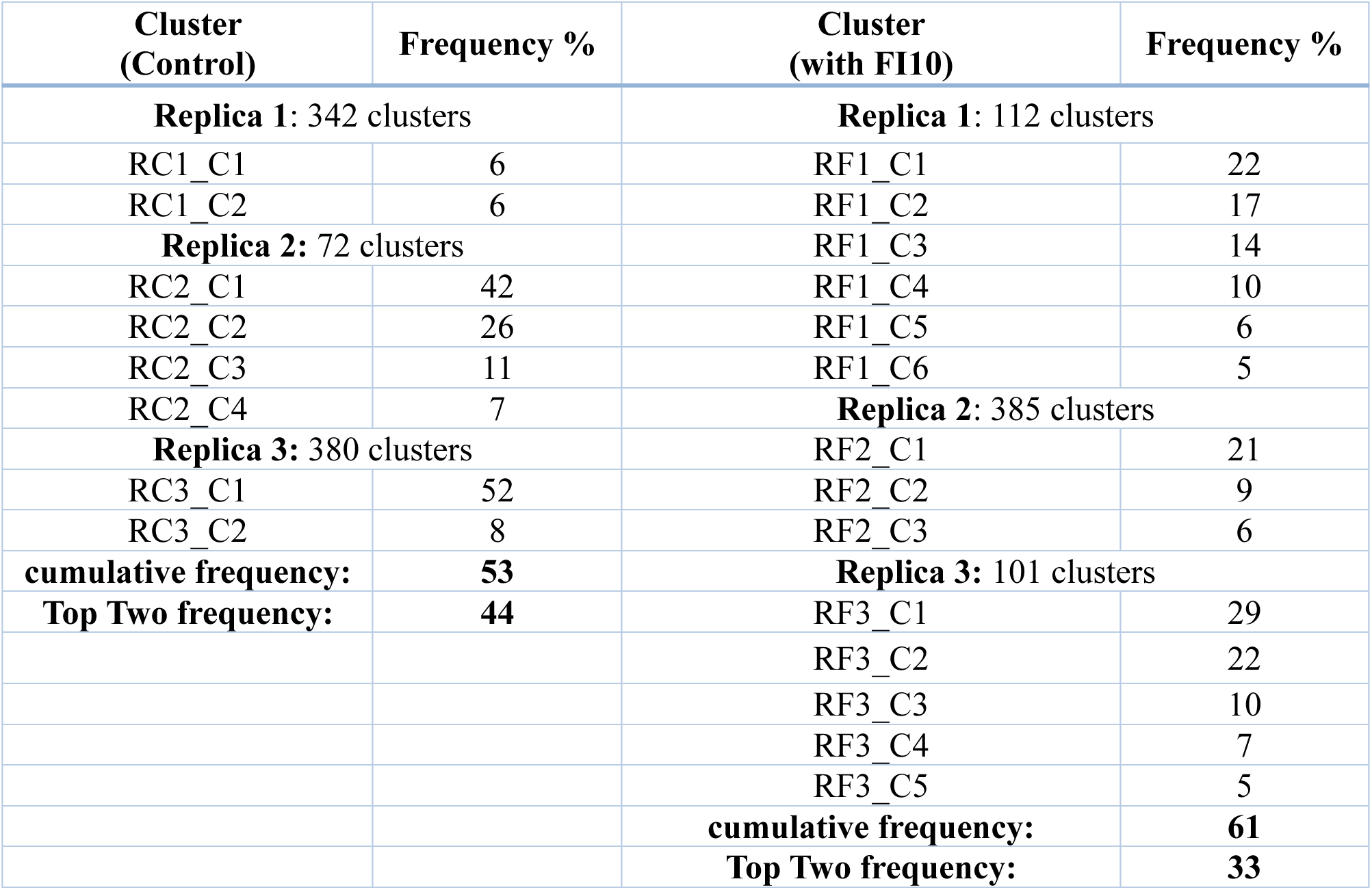
Frequency of the eight most populous clusters found in the control simulations, and the corresponding 14 clusters found in the simulations where FI10 interacts with Aβ_1-42_ monomer. Shown are also for both cases the total number of clusters and the cumulative frequency of the listed most populous clusters. Data are taken over the final 2.00 μs of each trajectory. Frequencies are rounded to the closest integer.

In a similar way, we find in the three trajectories following simulations of Aβ_1-42_ monomers interacting with FI10, six clusters containing more than 5% of the configurations in trajectory 1, three in trajectory 2 and five clusters in trajectory 3, with the frequencies of these 14 clusters, compromising together about 61 % of the 120000 snapshots in all three trajectories, again listed in **Table 2**, and their centroids shown in **Figure 6**.

**Figure 6.**
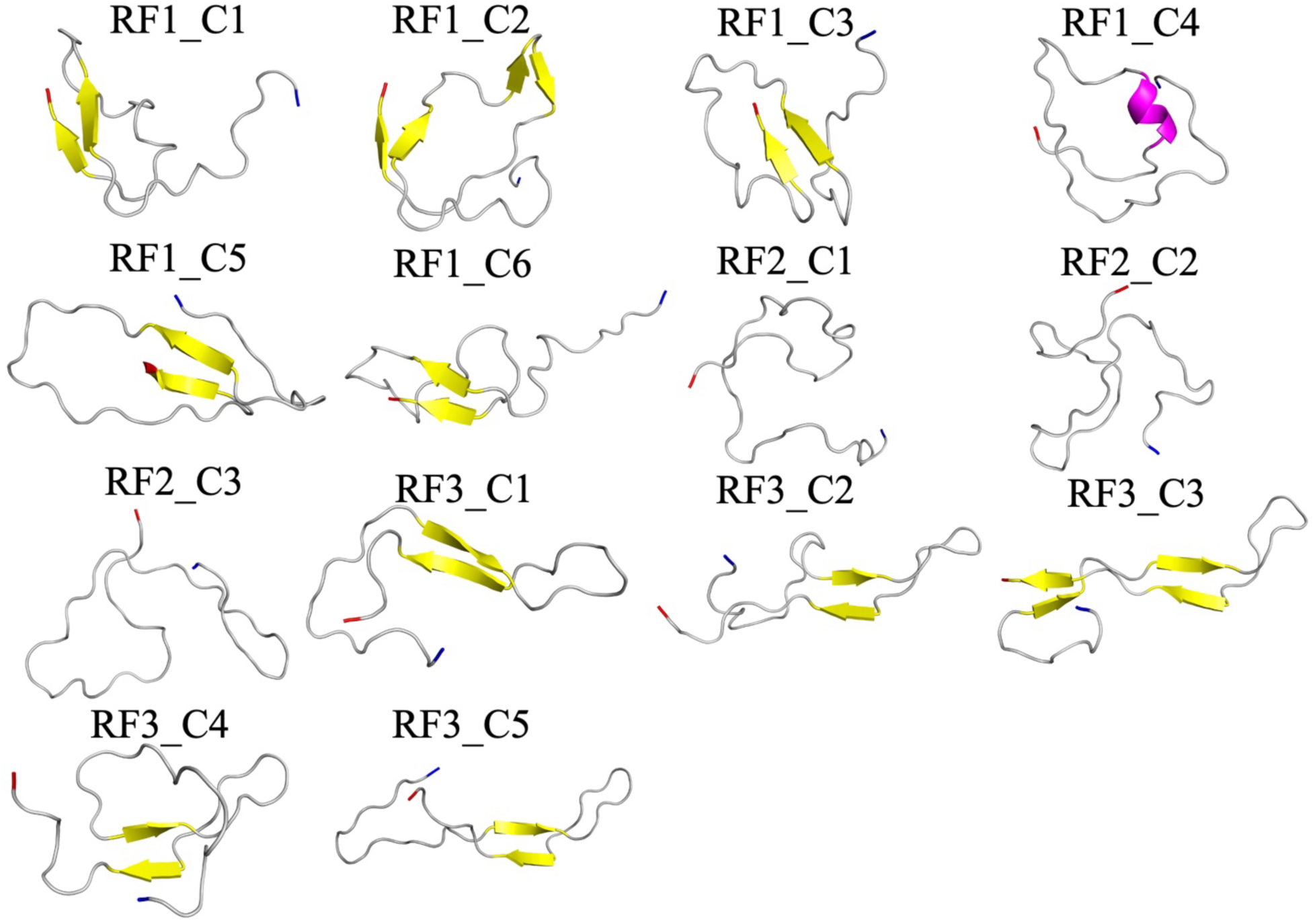
Ribbon representations of the centroid conformations in the 14 most populated clusters of configurations sampled over the final 2 μs in the three trajectories of simulations with FI10 present. The N- and C-terminals are colored in blue and red, respectively.

We notice that in presence of the FI10 viral fragment the monomer conformations cluster more tightly into 199 cluster (containing 61% of all conformations) than in the control where we find 265 clusters containing 53% of all conformations. Visual inspection of the cluster centroids suggests that the eight clusters from the control simulations can be characterized by the presence of certain transient secondary structure elements: β-strands containing three or more residues in the segment 16KLVFFAEDV24 (strand M1), 30AIIGLMV36 (strand C1), or 37GGVVIA42 (strand C2). This is confirmed by the corresponding plots of the strandness as function of residue number in **supplemental figure SF2**, where these segments appear as clearly identifiable regions. Note that the strand M1 corresponds to the hydrophobic segment M1 of **Figure 1**, and the strands C1 and C2 together form the hydrophobic C-terminal segment C of **Figure 1**. In six of the eight clusters (RC1_C1, RC1_C2, RC2_C1, RC2_C2, RC2_C3, RC2_C4) appears the strand M1 in conjunction with C1 or C2, see the frequencies and lifetimes of the three strand segments of the various clusters listed in **Table 3**, which also shows the respective averages taken over all control simulations. The correspondence of frequencies and similarities in lifetimes of M1 with C1 and C2, and the joined probabilities of M1⋂ C1 and M1⋂ C2 in **Supplemental Table ST1** indicate that presence of strand M1 is correlated with that of strand C2 and to a lesser degree C1 suggesting formation of a β-sheet between M1 and C2, which sometimes may also include C1.

**Table 3.** Frequency and life times of the strandness of the segments 10YEVHH14 (N1), 16KLVFFAEDV24 (M1), 30(AIIGLMV)36 (C1) and 37GGVVIA42 (C2) in all considered clusters, and averaged over all configurations sampled in the final 2 μs of each trajectory. Values are rounded to the closest integer. Standard deviation is given within the brackets.

|  | Segment |  |  |  |  |  |  |  |
| --- | --- | --- | --- | --- | --- | --- | --- | --- |
|  | N1 |  | M1 |  | C1 |  | C2 |  |
|  | Frequency % | Lifetime (ps) | Frequency % | Lifetime (ps) | Frequency % | Lifetime (ps) | Frequency % | Lifetime (ps) |
| Control | 0(0) | 104 (139) | 44 (36) | 592 (169) | 29 (22) | 541 (144) | 31 (28) | 539 (66) |
| with FI10 | 14 (17) | 347 (402) | 55 (44) | 1214 (1403) | 35 (56) | 1188 (1671) | 31 (29) | 553 (212) |
| Clusters |  |  |  |  |  |  |  |  |
| RC1_C1 | 0 | 50 | 37 | 266 | 14 | 151 | 36 | 267 |
| RC1_C2 | 1 | 50 | 88 | 231 | 36 | 157 | 86 | 228 |
| RC2_C1 | 0 | 0 | 85 | 527 | 81 | 605 | 80 | 427 |
| RC2_C2 | 0 | 0 | 66 | 747 | 0 | 0 | 11 | 470 |
| RC2_C3 | 0 | 0 | 88 | 415 | 74 | 308 | 86 | 411 |
| RC2_C4 | 0 | 0 | 86 | 298 | 44 | 208 | 84 | 305 |
| RC3_C1 | 0 | 0 | 1 | 123 | 1 | 155 | 8 | 384 |
| RC3_C2 | 0 | 50 | 2 | 63 | 1 | 75 | 0 | 0 |
| RF1_C1 | 52 | 235 | 62 | 406 | 8 | 671 | 74 | 389 |
| RF1_C2 | 15 | 164 | 66 | 308 | 19 | 357 | 69 | 340 |
| RF1_C3 | 65 | 337 | 76 | 996 | 0 | 0 | 80 | 902 |
| RF1_C4 | 33 | 165 | 55 | 226 | 0 | 0 | 71 | 236 |
| RF1_C5 | 3 | 142 | 0 | 0 | 0 | 0 | 73 | 707 |
| RF1_C6 | 23 | 128 | 57 | 182 | 0 | 0 | 63 | 177 |
| RF2_C1 | 0 | 0 | 23 | 154 | 0 | 100 | 23 | 154 |
| RF2_C2 | 0 | 0 | 0 | 0 | 0 | 0 | 0 | 0 |
| RF2_C3 | 0 | 0 | 49 | 458 | 0 | 0 | 38 | 388 |
| RF3_C1 | 6 | 307 | 100 | 1546 | 100 | 1555 | 9 | 330 |
| RF3_C2 | 0 | 0 | 99 | 2276 | 100 | 2786 | 1 | 80 |
| RF3_C3 | 38 | 380 | 100 | 346 | 100 | 346 | 40 | 363 |
| RF3_C4 | 0 | 0 | 100 | 3408 | 100 | 3882 | 0 | 0 |
| RF3_C5 | 7 | 157 | 100 | 562 | 100 | 562 | 21 | 258 |

Similarly, in the simulations where FI10 is present, one can also characterize the clusters by presence of transient secondary structure elements. However, besides the β-strands M1, C1, and C2 we also find now an N-terminal β-strand N1 centered around residues 10YEVHH14 and an α-helix made of residues A30-L34. Note, however, that, averaged over the three trajectories, less than 3% of the configurations contain transient helices while about 15% -20% of conformations have the strand-like segments N1, M1, C1, or C2. Similar to the control simulations, M1 seems to form in six clusters (RF2_C1, RF2_C3, RF3_C1, RF3_C2, RF3_C4, RF3_C5) a β-sheet with either C1 or C2, see the frequencies and life times of the four strand segments in **Table 3**, and the corresponding joined probabilities M1⋂ C1, N1⋂ C1, M1⋂ C2 and N1 ⋂ C2 in **Supplemental Table ST1**. The values in the two tables indicate for six other cluster (RF1_C1, RF1_C2, RF1_C3, RF1_C4, RF1_C6, RF3_C3) the formation of a β-sheet between N1 and C1 or C2, in addition to a sheet between M1 and C1 or C2. Note that the corresponding plots of the strandness as function of residue number for the various clusters in **Supplemental Figure SF3** indicate that presence of the viral fragment FI10 seems to shift the preference of residues to be strand-like toward the N-terminus of the segment M1 (residues 16 -24), so that sometimes N1 and M1 merge. This corresponds to a broad region of residues 10-20 to which FI10 has high binding affinity, see **Figure 3b**. At the same time, FI10 also seems to increase strandness of the C1 segment which again also corresponds to a region of high binding affinity. Life times of the M1 and C1 strands are almost double than in the control (see **Table 3**) while that of the C2 strand does not change. However, binding of M1 to C1 is often associated with that of N1 to C2, and both have comparable life times. Hence, presence of FI10 seems to lengthen the strands that interact with each other.

We remark that several studies have claimed that the central hydrophobic core composed of the M1 segment has a strong preference to form β strands and is highly amyloidogenic,^46–48^ and a fluorescence study has shown that the C2 segment composed of residues 37-42 exhibits strong fibrillation effect by stabilizing β-hairpin structures.^49^ Hence, while due to differences in simulation protocol, secondary structure determining method, and using different starting structures and force fields the frequencies of strands M1 and C1 in our study are higher than in an earlier computational study (where these strands were observed with frequencies of ∼6-8% and ∼8-14%, respectively),^50^ the increase in both the frequency and lifetime of β-strands in presence of FI10 still supports our hypothesis that the viral protein fragment facilitates Aβ_1-42_ aggregation.

What keeps the strands connected and stabilized? In **Table 4** we list the average number of hydrophobic contacts between the segments when in strand-like conformation, and compare it with corresponding numbers for arbitrary residues. On average, about 30 (8) such contacts are found over the whole trajectory in the control simulations. The maximum of possible contacts is 276 (24*23/2), i.e., the density of hydrophobic contacts is on average 0.11. On the other hand, if M1 and C1 are in a strand conformation, there are about ten (of 42 possible) hydrophobic contacts between the two strands, corresponding to a density of 10/42 i.e., 0.24. Hence, hydrophobic contacts between residues in these two segments are found 2.2 times more frequently than on average. If M1 and C2 are in a strand conformation one finds on average 16 of 36 possible hydrophobic contacts between the two segments, leading to a density of around 0.44, i.e., such contacts are found 4 times more frequently than on average. In the simulations where FI10 interacts with the Aβ_1-42_ monomer, we observe on average 27 (5) contacts between the 24 hydrophobic residues, i.e., the density of such contacts is similar to ones in the control simulations. However, we find now almost the double number (≈19) of contacts between M1 and C1, and about half the number (≈9) of contacts between M1 and C2. Added to these numbers are one additional hydrophobic contact between the segments N1 and C1 and three contacts between segments N1 and C2. This increase in the number of hydrophobic contacts and the relative shift between the segments agrees with our previous observation that presence of FI10 lengthen the strands that interact with each other.

**Table 4.**
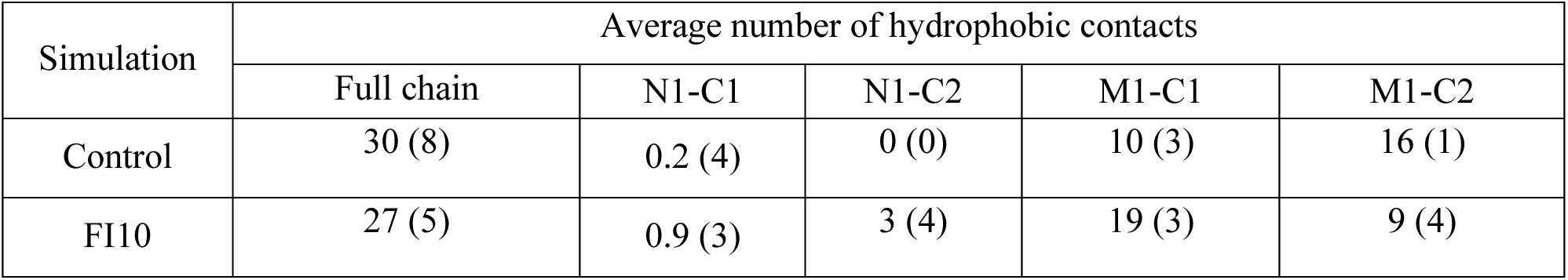
Number of hydrophobic contacts between residues in the Aβ_1-42_ monomer, for the full chain, and between the segments N1, M1, C1, and C2 when in a strand conformation. Data are averaged over all configurations sampled in the final 2.00 μs of each trajectory. Standard deviation is given within the brackets.

We remark that we did not observe hydrogen bonds between segments N1 or M1 and C1 and C2 in our simulations, independent of presence or absence of the viral protein fragment FI10. However, the transient β-sheets and strands seen in our monomer simulations may also be stabilized by salt bridges between charged residues. Once formed, these salt bridges would restrict the conformational space of the segments N1, M1, C1 and C2, encouraging in this way formation of the observed secondary structure elements, and that process may be modulated by presence of the viral spike protein fragment FI10. We have therefore also looked into the effect of FI10 on salt bridge formation and the correlation between observed salt bridges and the transient β-sheets and strands seen in our monomer simulations. When restricting ourselves to salt bridges that appear with more than 5% in at least one trajectory, we find three such salt bridges: E11-H13, E22-K28 and D23-K28. The average frequency and life times of these salt bridges are listed in **Table 5**.

**Table 5.** Average frequency of appearance of salt bridges and lifetime of salt bridges. Shown are values calculated over all respective trajectories, and such restricted to cases where the segments 10YEVHH14 (N1), 16KLVFFAEDV24 (M1), 30(AIIGLMV)36 (C1) and 37GGVVIA42 (C2) are in a strand conformation. Data are averaged over all configurations sampled in the final 2.00 μs of each trajectory. Standard deviation is given within the brackets. Values are rounded to the closest integer.

| Frequency % |  |  |  |  |  |  |
| --- | --- | --- | --- | --- | --- | --- |
|  | E11-H13 |  | E22-K28 |  | D23-K28 |  |
|  | Control | FI10 | Control | FI10 | Control | FI10 |
| Full chain | 4 (3) | 16 (12) | 23 (25) | 6 (5) | 4 (4) | 29 (38) |
| N1 | 0 (0) | 30 (15) | 29 (26) | 2 (1) | 1 (2) | 24 (28) |
| M1 | 5 (3) | 16 (12) | 10 (12) | 3 (3) | 2 (2) | 47 (32) |
| C1 | 7 (3) | 8 (6) | 8 (9) | 1 (3) | 3 (3) | 68 (17) |
| C2 | 5 (2) | 24 (11) | 4 (13) | 7 (4) | 0 (0) | 16 (23) |
| N1∩C1 | 0 (0) | 7 (12) | 16 (37) | 2 (3) | 0 (0) | 63 (28) |
| N1∩C2 | 0 (0) | 29 (15) | 15 (35) | 2 (1) | 2 (4) | 24 (28) |
| M1∩C1 | 6 (3) | 8 (4) | 8 (10) | 1 (3) | 3 (3) | 70 (13) |
| M1∩C2 | 5 (2) | 26 (12) | 1 (1) | 6 (3) | 0 (0) | 18 (25) |
| Life time (ps) |  |  |  |  |  |  |
| Full chain | 105 (5) | 115 (18) | 614 (282) | 388 (199) | 754 (381) | 928 (502) |
| N1 | 0 (0) | 108 (20) | 163 (53) | 130 (14) | 0 (0) | 268 (159) |
| M1 | 103 (12) | 103 (2) | 322 (9) | 128 (44) | 472 (248) | 789 (381) |
| C1 | 103 (7) | 102 (9) | 152 (167) | 96 (12) | 327 (107) | 1062 (267) |
| C2 | 93 (21) | 105 (10) | 193 (40) | 224 (49) | 495 (251) | 349 (94) |
| N1∩C1 | 0 (0) | 137 (29) | 8 (18) | 106 (14) | 0 (0) | 500 (223) |
| N1∩C2 | 0 (0) | 109 (22) | 0 (0) | 128 (15) | 0 (0) | 284 (177) |
| M1∩C1 | 96 (8) | 103 (3) | 157 (165) | 94 (6) | 312 (118) | 1060 (192) |
| M1∩C2 | 97 (6) | 106 (10) | 181 (33) | 165 (37) | 514 (242) | 345 (101) |

Our data in **Table 3** and **Supplemental Table ST1** indicate that the four observed strands form transiently even in the control, but that the frequency of strands and their pairing is higher in presence of FI10. On the other hand, when considering the frequencies of the three salt bridges calculated for each system over all three trajectories and listed in **Table 5**, we see that the salt bridge E22-K28 is more common in the control than in presence of FI10, while the opposite is true for the salt bridge between E11 and H13, and the one between D23 and K28. The reduced frequency of the salt bridge E22-K28 and its halved lifetime in presence of FI10 correlates with the binding pattern of the of the viral protein fragment to the Aβ_1-42_ monomer in **Figure 3**, i.e., it appears that binding of FI10 to the monomers interferes with formation of the salt bridge. We also note that while the standard deviations in our frequencies are large, even in the control seems the existence of this salt bridge to be anticorrelated with the strandness in the segments N1, M1, C1 and C2.

On the other hand, frequency and lifetime of the D23-K28 salt bridge are not only higher in presence of FI10 than in the control, but also larger when the sheets between N1 and C1 or between M1 and C1 are formed, and similar for the sheets between N1and C2 and between M1 and C2. These results are consistent with many experimental and theoretical studies showing that the intramolecular D23-K28 salt bridge facilitates the oligomer stability and fibrilization process of Aβ_1-40_ and Aβ_1-42_ peptides.^50–54^ Hence, the enhanced lifetime and frequency of D23-K28 salt bridge indicates that presence of FI10 could facilitate the aggregation of Aβ_1-42_ monomers by encouraging formation or stabilizing this salt bridge. Note, however, that the probability of finding the salt bridge D23-K28 is higher under the condition that both N1 and C1 (or M1 and C1) are formed than for the condition that N1 or M1 are formed independently of whether C1 forms a strand. This indicates that presence of this salt bridge depends on formation of a sheet between the segments N1 and C1, or between segments M1 and C1, i.e., the salt bridge D23-K28 is formed after the sheet between segments M1and C1, not causing it. Note also that the sheet between segments M1 and C1 is found in control simulations with a probability of 18% and a life time of 750ps, while the corresponding frequency in presence of FI10 is 34% with a lifetime of 1000ps. In presence of the salt bridge D23-K28 increase frequency and life time in the control simulations to 29% and 750ps, while in the FI10 simulations the life time stays unchanged and the frequency increases to about 82%, see **Supplemental Table ST2**. Hence, it appears that the sheet formation is not caused by the D23-K28 salt bridge but rather by direct interaction with the viral fragment or induced hydrophobic contacts.

A similar picture is seen for the salt bridge E11-H13 which appears in the control simulations with about 4%, but in presence of FI10 with around 16%, see **Table 5**. In the presence of FI10 increases this frequency to about 30% if the strand N1 is formed, a value that is similar to the one in presence of sheets between segments N1and C2 or M1 and C2. Note that little differences are seen in the lifetimes of the salt bridge, neither changing with presence of FI10 nor depending on the presence of strands. Considering the opposite relationship, i.e., the frequency of strandness in the segments as a function of presence of the S11-H13 salt bridge (**Supplementary Table ST2**), we see that even in the control presence of the salt bridge increase the strandness of the segments M1 leading to a higher frequency for the sheets between M1 and C1 or M1 and C2. Correspondingly, we see in presence of FI10 that existence of the E11-H13 salt bride implies a higher frequency for the sheets between segments N1 and C2, and M1 and C2. Hence, formation of the N1 strand is likely resulting from the binding of FI10 to residues Y10 to F20 (segment N1 and parts of M1) that also increase the probability to form the E11-H13 salt bridge. Once this salt bridge is formed, it leads to the observed pattern of residues 10-20 interacting with residues 30-40. Hence, unlike for the D23-K28 salt bridge, we find that presence of FI10 increases the frequency of the E11-H13 salt bridge which in turn then eases interaction between the N-terminal and C-terminal segments.

### Fibril simulations

Aβ_1-42_ amyloids form by association of monomers, but the propensity of amyloids does not only depend on the ensemble of monomers (containing more or less of aggregation-prone conformations) but also on the stability of the final product of these associations, the experimentally observed fibrils. Hence, viral protein fragments can enhance Aβ-amyloid formation both by shifting the monomer ensemble toward more aggregation prone conformations and by altering the stability of the fibrils. In previous work we have shown such stabilization by the FI10 fragment for αS fibrils, and found that the effect is differential, i.e., depends on the specific fibril structure.^13^ Aβ-fibrils are characterized by a large polymorphism with the structural differences correlating with the severity of disease symptoms.^20,55^ Hence, differential stabilization of Aβ_1-42_-fibrils could potentially hasten the outbreak and/or pathogenesis of Alzheimer’s disease. For this reason, we have considered in this work also the effect of FI10 fragments on two distinct fibril models. The first one, having the PDB-ID 7Q4B^56^, is derived from the brains of deceased Alzheimer disease patients, while the second one (with PDB-ID 5KK3^57^) is a synthetic fibril, i.e., the aggregation happened *in vitro* and not the brain environment. Choice of the two models therefore allows us to probe whether FI10 alters the stability of Aβ_1-42_-fibrils, and whether this effect depends on the fibril structure and therefore may modulate the pathogenesis of Alzheimer’s disease. Cartoons of the two fibril models are a shown in **Figure 1e**-**1f**.

Patient-derived fibrils of type 1, the ones considered in this study, are present in the brains of sporadic Alzheimer’s disease patients, while the type 2 fibrils (not discussed in this study) are found in the brains of familial Alzheimer’s disease patients.^56^ The fibril is composed of two S-shaped protofilaments which each contain five β-strands per chain. The protofibrils bind through hydrophobic interactions involving the side chains of L34, V36, V39, I41 residues of S-shaped domain and Y10, V12, Q15, and L17 of N terminal arm.^56^ The synthetic fibril model considered in this study is also composed of two S-shaped protofibril that each contains in every chain four β-strands between residues 16 and 42.^57^ Some hydrophobic interactions including F19-I32, F19-A30, F20-A24, V24-G29, G29-I41, I31-V36, and G33-V36 play a vital role in determining the fold of monomer structure in the synthetic fibril. Intermolecular interactions L17-M35 and Q15-M35 have been reported as important interactions between the two protofibrils.^57^

We start our investigation by considering for each system the root-mean-square deviation (RMSD) to the respective fibril conformation as function of time. This quantity is calculated for backbone atoms only ignoring flexible N-terminal residues (the first eight residues of the patient-derived fibril and the first ten residues of synthetic fibril), and in **Figure 7a**-**7b** we show for each system its average over three trajectories. This quantity describes the overall change in the fibril structure along the trajectories, while the change in the structure of individual chains is described by the chain RMSD, which is the average over the RMSD calculated separately for each chain. We show the chain RMSD in the insets. For both fibril models we observe that presence of FI10 leads to a slightly higher chain rmsd, i.e., a de-stabilization of the chain conformations. However, the effect of FI10 on the overall RMSD depends strongly on the fibril model. Little effect is seen for the synthetic fibril (5KK3) while the patient-derived fibril (7Q4B) is de-stabilized in presence of FI10. A consequence of these changes are seen in the solvent accessible surface area (SASA) also shown in **Table 6**, which in the patient-derived fibril only marginal changes with time and is not affected by presence of FI10, but is in the synthetic fibril with 264 (5) nm^2^ about 23 nm^2^ larger in presence of FI10 than in the control (241 (13) nm^2^), with 13 nm^2^ resulting from addition of hydrophobic surface area. We expect that the differences in the overall RMSD and the SASA indicate different re-arrangement of the Aβ_1-42_ in the two fibril geometries over the course of the trajectories caused by the presence of FI10 peptides. As both models are built out of two protofibrils, each made of five layers, changes in the arrangement of the chains can come from loosing or tightening of the layers in each protofibril, or from loosening or tightening of the two protofibrils. The first effect can be quantified by the change in the number of stacking contacts between layers, while the second effect described by the change of packing contacts between the protofibrils.

**Figure 7.**
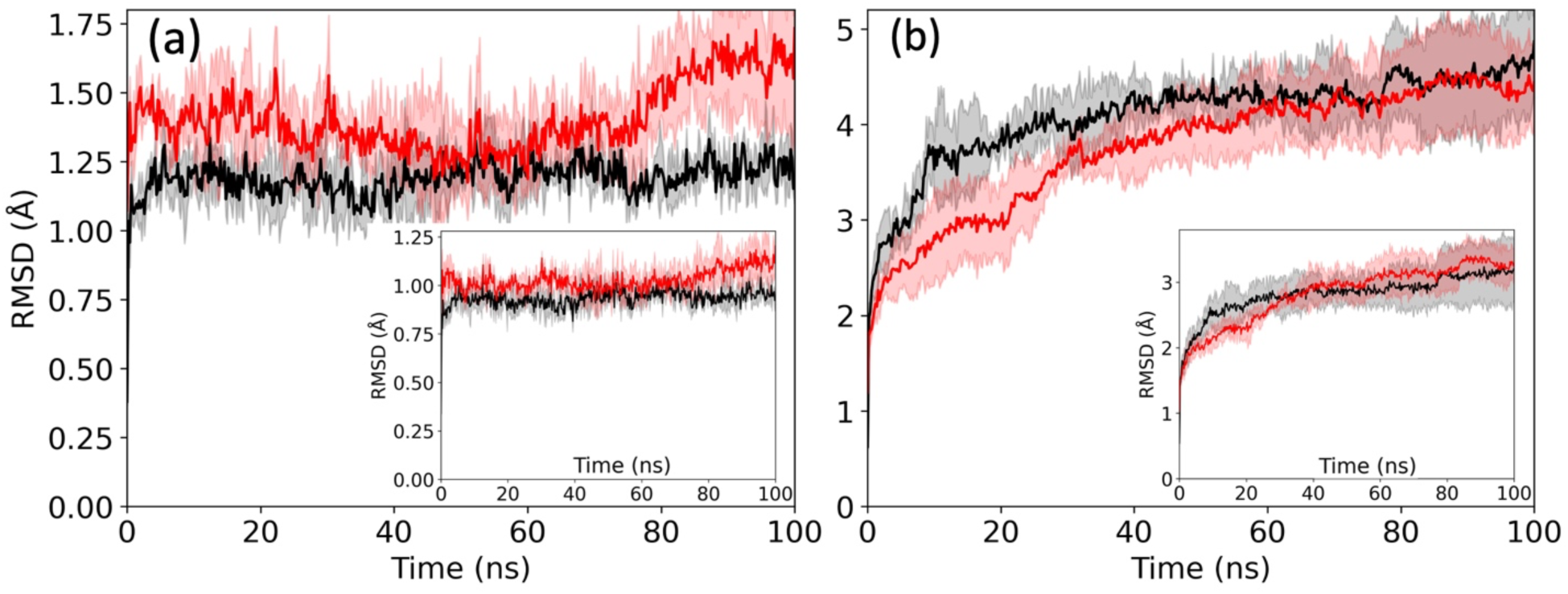
Root-mean-square deviation (RMSD) of the fibril conformations for the patient-derived fibril 7Q4B (a) and the synthetic fibril 5KK3 (b) to the respective fibril model (the start configuration) as function of time. Values for the control are drawn in black and the one from simulations with FI10 present in red. The insets show the chain RMSD, i.e., where the RMSD is calculated separately for each, and averaged over all ten chains of the system.

**Table 6.** Average number of total contacts, intrachain contacts, staggering contacts and packing contacts, and solvent accessible surface area (SASA), calculated over the last 50 ns of all three trajectories for each system. The initial values (measured at 0.2 ns to account for steric clashes observed at t=0)) for each quantity are also given. Note that we do not consider contacts involving the N-terminal residues as the N-terminal segment is flexible (see main text for details). Native contacts are such also seen in the respective fibril models and long-living packing contacts defined in the main text.

|  | Patient derived fibril (7Q4B) |  |  |  | Synthetic fibril (5KK3) |  |  |  |
| --- | --- | --- | --- | --- | --- | --- | --- | --- |
|  | Control |  | FI10 |  | Control |  | FI10 |  |
|  | Initial | Last 50 ns | Initial | Last 50 ns | Initial | Last 50 ns | Initial | Last 50 ns |
| Intrachain contacts | 16.5 (4) | 16.9 (2) | 15 (1) | 17.0 (3) | 13.9 (5) | 14 (1) | 13.2 (7) | 12 (1) |
| Stacking contacts | 170 (2) | 169 (1) | 168 (2) | 166 (1) | 148 (6) | 139 (7) | 143 (2) | 142 (5) |
| Native Stacking Contacts | 158 (1) | 146 (1) | 150 (0) | 140 (1) | 129 (4) | 100 (7) | 122 (3) | 98 (8) |
| Packing contacts | 99 (6) | 120(9) | 89 (5) | 119 (3) | 108(5) | 178 (17) | 106 (9) | 140(23) |
| Native Packing Contacts | 86 (7) | 71(3) | 69 (3) | 55 (3) | 87 (10) | 47 (3) | 80 (5) | 46(12) |
| Long-living Packing Contacts | 99(6) | 110(9) | 89(5) | 98(6) | 108(5) | 108(18) | 106(9) | 98(12) |
| Packing Distance (Å) | 11.7 (2) | 11.3 (1) | 12.0 (1) | 11.5 (3) | 9.3 (2) | 9.6 (7) | 9.8 (2) | 9.2(4) |
| SASA (nm<sup>2</sup>) | 203 (4) | 205 (2) | 210 (1) | 212 (2) | 284 (7) | 241 (13) | 275(4) | 264 (5) |
| hydrophobic<br>SASA<br>(nm<sup>2</sup>) | 93 (2) | 94 (1) | 100 (0) | 98 (1) | 144 (3) | 120 (8) | 142 (2) | 133 (3) |
| hydrophilic<br>SASA<br>(nm<sup>2</sup>) | 110 (2) | 111 (2) | 110 (1) | 114 (2) | 140 (4) | 121 (5) | 133 (2) | 131 (2) |

In **Figure 8a**-**8b** we show the time evolution of the number of staggering contacts (averaged over all three trajectories). Note that we do not consider contacts involving the flexible N-terminal segment added by us and missing in the fibril models. For an easier comparison we have normalized the plots such that it is unity at 0.2 ns - a value chosen to account for steric clashes at t=0 ns. In both the control and in presence of FI10 we see little change along the trajectories for the patient-derived fibril, and a decrease of in absolute numbers of about ten contacts in the control for the synthetic fibril. When the values are averaged over the final 50 ns, we find for both fibrils similar differences between control and FI10 simulations, indicating that conformations in the control simulations are stabilized by about 2-3 staggering contacts more than in the simulations where FI10 is present, see **Table 6**. Note that the contacts seen at the end of the trajectories are not necessarily the ones present in the respective fibril models. The number of these native stagging contacts decreases in all cases, see **Figure 8c**-**8d**, but stays for both fibril models slightly higher in presence of FI10, see also the averages over the last 50 ns displayed in **Table 6**. Hence, presence of FI10 seems to stabilize the original binding between the layers in both the patient-derived fibril and in the synthetic fibril. However, newly formed contacts only partially compensate for the loss of native staggering contacts, and fewer are formed in presence of FI10, leading in both fibril models to an effective weakening through presence of FI10 by about 2-3 contacts.

**Figure 8.**
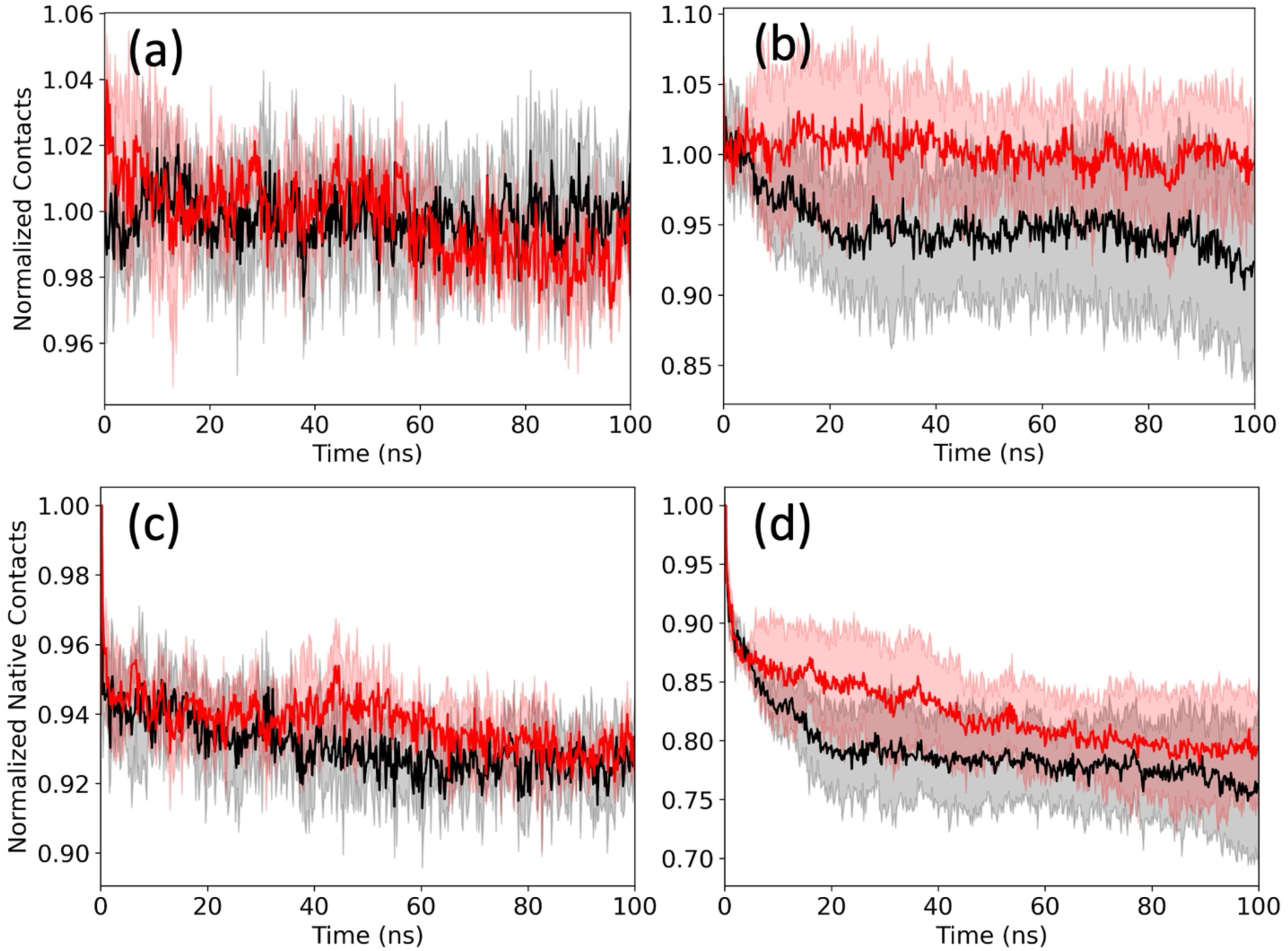
Average number of normalized stacking contacts between residues in neighboring layers of the fibril conformations for the patient-derived fibril 7Q4B (a) and the synthetic fibril 5KK3 (b) as function of time. Values for the control are drawn in black and the one from simulations with FI10 present in red. In (c) and (d) we show the corresponding figures for native contacts, that is contacts which are seen in the respective fibril models.

The situation is more diverse for the time evolution of normalized packing contacts and native packing contacts shown in **Figure 9**. For the patient-derived fibril decreases the number of native packing contacts both in the control and in presence of FI10, but is lower in presence of FI10. However, the loss of native packing contacts is more than compensated by formation of new packing contacts, and for the control the absolute number plateaus quickly while it grows in presence of FI10. Final values, averaged over the last 50 ns are again shown in **Table 6** indicating that the presence of FI10 30 additional contacts are formed while only 21 in the control. For the synthetic fibril we also observe a decrease in the number of native packing contacts, with the loss similar in presence and absence of FI10. Averaged over the last 50 ns we find 178 (17) contacts in the control simulations, but only 140 (23) contacts in presence of FI10, see **Table 6**. We remark that we did not see similar differences in the number of intrachain contacts whose time evolution is shown in **Supplementary Figure SF4,** and for which average values for the last 50 ns are also listed in **Table 6**. Here, the numbers of intra chain contacts is for the patient-derived 7Q4B fibril with. about 17 contacts essentially the same in presence and absence of FI10, and is only marginally smaller in presence of FI10 (12 (1) versus 14 (1) for the control) for the synthetic 5KK3 fibril, confirming our assumption that presence of FI10 alters the arrangement of the Aβ_1-42_ - chains, not their structure.

**Figure 9.**
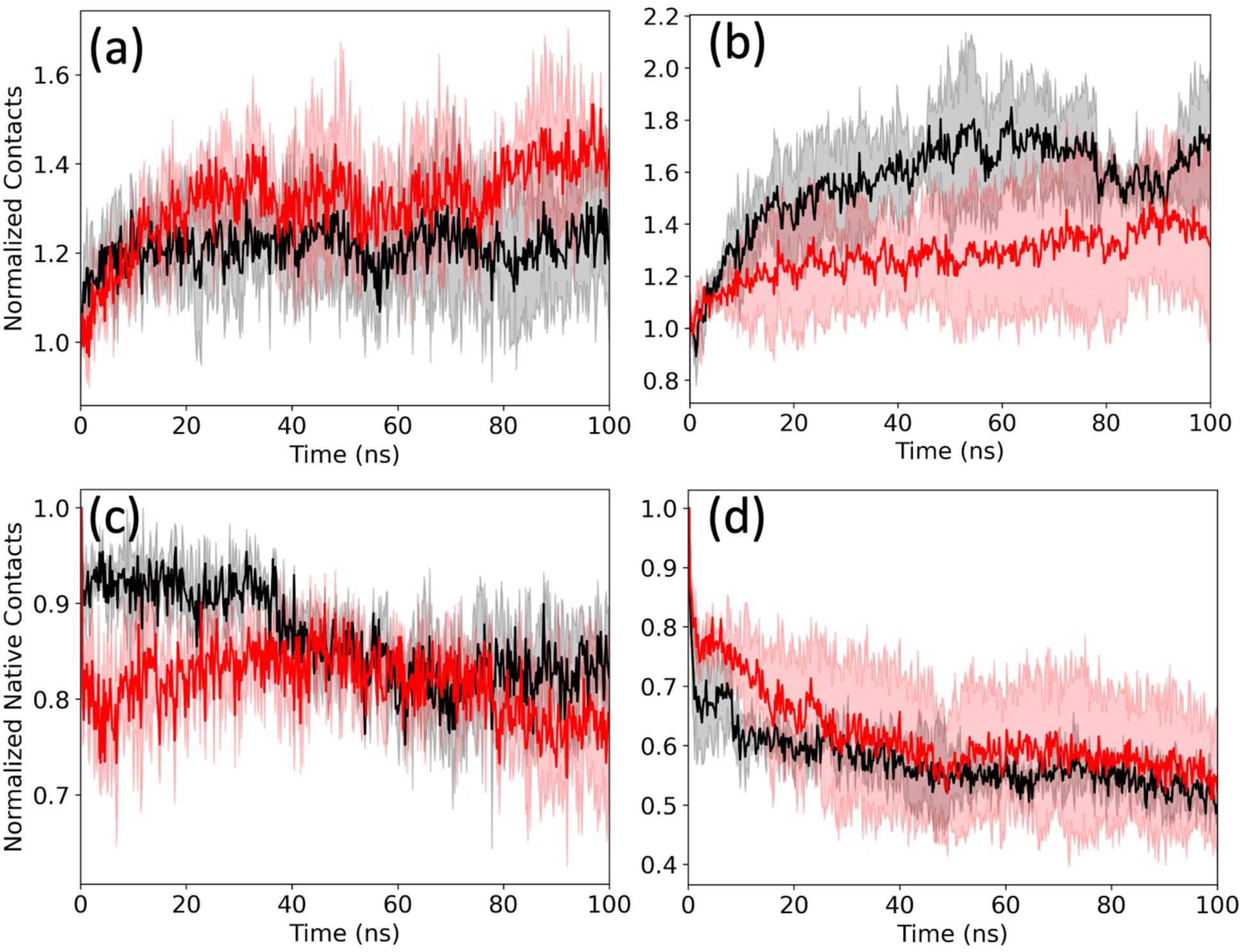
Average number of packing contacts between residues in opposite protofibrils of the fibril conformations for the patient-derived fibril 7Q4B (a) and the synthetic fibril 5KK3 (b) as function of time. Values for the control are drawn in black and the one from simulations with FI10 present in red. For better comparison are the values normalized to unity for the conformation observed at start. In (c) and (d) we show the corresponding figures for native contacts, that is contacts which are seen in respective fibril models and the start conformations.

This change in packing contacts is correlated with that in the packing distance, that is the distance between the two protofibrils. We define packing distance as the distance between center of masses of interfacial residues of two protofibrils. For the patient-derived fibril, we considered Y10, V12, Q15, L17, L34, M35, V39, I41 as interfacial residues, while the residues H13, H14, Q15, K16, G33, L34, M35, V36, G37, G38, V39 were considered as interfacial residues of synthetic fibril. For the patient-derived fibril this packing distance decreases in the control by about 0.4 Å and 0.5 Å in presence of FI10; i.e., the two protofibril change their relative positions (and therefore the contacts between residues) but their distance to each other is little changed and slightly more reduced in presence of FI110. The stabilizing effect on packing by FI10 is also seen for the synthetic fibril where the distance decreases in presence of FI10 but increases in the control. Hence, it appears that FI10 slightly destabilize the stacking of the Aβ_1-42_-chains in a way that does not depend on the specific fibril type, but has a stronger effect on the packing of protofibrils in a way that depends on the fibril geometry.

For the patient-derived fibril are the main packing contacts in the PDB-structures such between V12-V39, L17-V36, L34-V36, V36-L34, V36-L17 and V39-V12, with the residues on different proto fibrils, and contacts either with residues of the same layer or shifted by one layer. When interacting with FI10 the number of such contacts is reduced from about 44 to 41. However, these contacts survive in both cases throughout the simulations, and the observed changes in the number and kind of packing contacts results from re-arrangement of other packing contacts such as Q15-V39 or F19-V36, with new contacts often between residues in different layers in the protofibrils, and their number larger in presence of FI10 (23 as opposed to 14 in the control), leading to the decrease in packing distance seen in both control and presence of FI10. On the other hand, in the synthetic fibril are the dominating contacts L17-M35, L34-M35, M35-L34 and M35-L17, which however decrease from about 30 for both cases at start to 23 in the control and only 19 in presence of FI10. Interestingly, the loss of these contacts is mostly for such contact between residues on different layers (about 7), and is for residue pairs located on the same layer smaller than in the control (3 as opposed to 8 in the control).

Note, that as already reported above, the total number of packing contacts increases in both systems. However, most of these newly formed contacts are transient. This can be seen if we measure instead of the contacts the number of pairs where the *average distance* between the residues is smaller than the cut-off distance of 4.5Å used in our definition of a contact. Such pairs correspond to long-living packing contacts, while our previous definition also accounts also for short-live and transient contacts. At start we find for the synthetic fibril in the control 108 (5) of such pairs, and in presence of FI10 106 (9) pairs, but in the last 50 ns 108 (18) pairs in the control and only 98 (12) in the trajectories where FI10 is present. Hence, the number of long-living contacts decreases as does the number of native contacts (see above), and the newly formed non-native contacts are therefore short-lived transitory contacts that do not lead to a decrease in the packing distance, i.e., a stronger interaction between the two protofibrils. This is different in the patient-derived fibril where the number long-living packing contacts also increases for the control simulations by about 11, and by about 9 in presence of FI10. These additional contacts lead to a stronger interaction between the two protofibrils in the presence of FI10, causing a reduction of packing distance and therefore stabilizing the fibril.

The observed differences may result from the disparate binding pattern of FI10 two the two fibrils geometries, see **Figure 10**. For the patient-derived fibril binds FI10 with a free energy of about - 90 (8) kJ/mol to the fibril. **Figure 10a** shows that binding of FI10 is preferably to the residues 15-23 of the hydrophobic segment M1 (on average 51%) and residues 30-41 of hydrophobic segments C1 and C2 (on average 33% to the C1 segments, with the maximum of 44 % at L34, and on average with 20% to the C2 segment), with a binding probability of about 70% for the charged residues K16 and E22 or D23. FI10 binds with other parts of fibril with a much lower affinity of about 7%. On the other hand, the binding distribution is more indiscriminate for the synthetic fibril with in general higher binding probabilities (on average 35%), but no specific preference for a certain segment, see **Figure 10b**. When looking into the binding pattern of the FI10 residues, we find that the Aβ_1-42_ chains of the synthetic fibril also bind unspecific to FI10 residues (**Figure 10f**), but at higher binding frequencies than seen for binding to the patient-derived fibril where in addition the binding propensities slightly decrease from N- to C-terminal of FI10 (**Figure 10e**). This suggests that while on the synthetic fibril FI10 is with a free energy of -108 (9) kJ/mol more tightly bound than when on the patient-derived fibril, the peptide is on the synthetic fibril mobile and changes the residues on the chains with that it interacts, while on the patient-derived fibril it sits more localized. Hence, the lower RMSF and lower flexibility in the synthetic fibril in presence of FI10 (**Figure 10d**), while for the patient-derived fibril the interaction with FI10 leads to increased flexibility of the not interacting chain segments, especially the N-terminus (**Figure 10c**).

**Figure 10.**
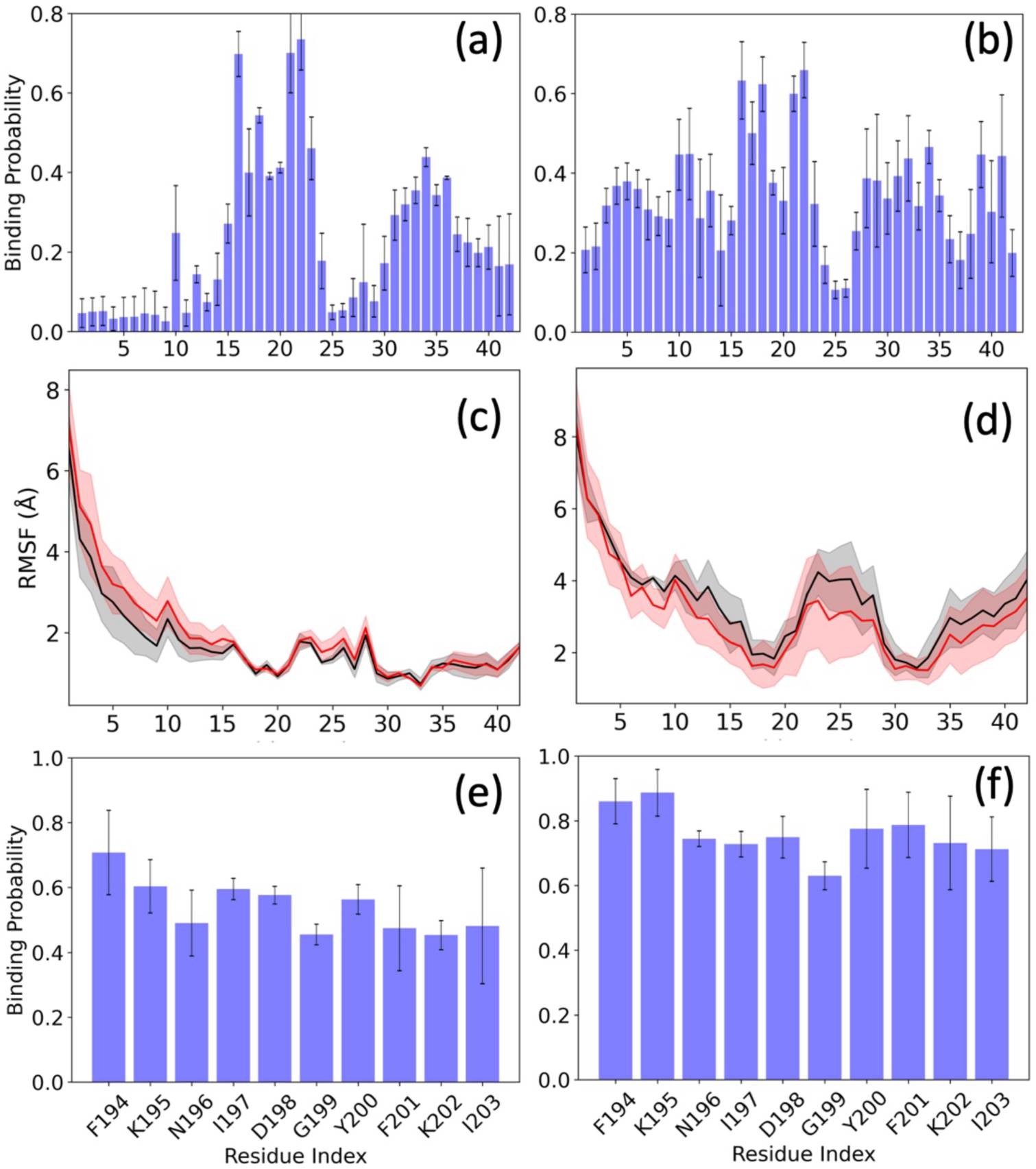
The residue-wise normalized binding probability of the FI10 segment for the Aβ_1-42_ chains is shown in (a) for the patient-derived fibril model 7Q4B and in (b) for the synthetic fibril model 5KK3, while the corresponding residue-wise mean square fluctuation (RMSF; in Å) obtained from Aβ_1-42_ fibril simulations in the absence (in black) and presence (in red) of the FI10 are shown in (c) and (d). In (e) we draw the complementary, and in (f) the binding probability of residues in the synthetic fibril model 5KK3 to FI10 residues. Data are averaged over the final 50ns of each trajectory. Shaded regions or error bars mark the standard deviation of the averages.

## Conclusions

In order to understand the effect of small viral protein fragments on the aggregation of Aβ_1-42_ peptides as a potential factor in the initiation and progression of Alzheimer’s disease, we have performed all-atom molecular dynamics simulations of Aβ_1-42_ monomer and fibril in presence of SARS-COV-2 Spike protein fragment 194FKNIDGYFKI203 (FI10). Our simulations indicate that FI10 shifts the ensemble of monomers to less compact conformations with a larger surface exposed to the solvent, stabilizing especially of the C-terminal segment of residues 30 to 42. While even in absence of FI10 strand-like segments 16KLVFFAEDV24 (strand M1), 30AIIGLMV36 (strand C1), or 37GGVVIA42 (strand C2) form, presence of FI10 increases their frequency and lifetimes and extends the segment M1 toward the N-terminus or adds a strand 10YEVHH14 (N1). The N- and C-terminal segments are stabilized by hydrophobic contacts between the segments that are higher in presence of FI10; and further stabilization is provided by salt bridges D23-K28 and E11-H13 forming subsequently. Hence, presence of FI10 will increase the propensity of Aβ_1-42_ monomers to aggregate. However, this increase is unspecific as the aggregation-prone strand segments appear not to favor a specific fibril polymorph and are found in many of the experimentally resolved fibril models, see, for instance, the fibril models with PDB IDs 7Q4B, 7Q4M, 5KK3, 8EZD.

As the equilibrium between monomers and amyloids can be also shifted by stabilizing the end product of this process, we have investigated in complementary simulations two distinct fibril fragments, the patient derived 7Q4B and the synthetic 5KK3, as both contain the above-described segments N1, M1, C1 and C2. Analyzing our trajectories, we find no change by FI10 on the chain conformations (for instance, in the number of intra-strand contacts), but on the relative arrangement of the chains in the fibrils. FI10 may slightly destabilize the stacking of the Aβ_1-42_-chains in a way that does not depend on the specific fibril type, but has a stronger effect on the packing of protofibrils (as seen in number of packing contacts, packing distance and the solvent exposed surface area) that depends on the fibril geometry. Especially, we find that presence of FI10 leads to a stronger packing of the proto fibrils in the patient-derived fibril, while this effect is only marginal for the synthetic fibril. Hence, while FI10 increases the aggregation propensity of the monomers without obvious preference for a specific outcome, it stabilizes the fully-formed fibrils differently depending on their geometry. We relate this differential stabilization to the more localized binding of FI10 to the patient-derived fibril than seen in the synthetic fibril where FI10 is binding diffusively.

Comparing our results for of Aβ_1-42_ monomers and fibrils with earlier work, we find as a common theme that FI10 and other SARS-COV-2 protein fragments can indeed increase amyloid formation of human proteins such as αS, SAA and amylin.^12,13,21,22^ As such small viral protein fragment are likely present *in vivo* following cleavage during infection-caused inflammation, amyloid diseases may be triggered in this way by SARS-COV-2 and likely other viral infection. Common seems to be in all cases that the viral protein fragments shift the ensemble of monomer conformations toward more aggregation-prone ones. Less clear is the effect on the end product, the fibrils often seen as a hallmark of various diseases. Here we find that the modulation of stability of the potentially cell-toxic fibrils depends on the geometry of the fibrils. In the case of Aβ_1-42_ it appears that fibril polymorphs are stabilized that are found in patients with Alzheimer’s disease. However, our study looked only into two of many experimentally resolved fibrils, and more work is needed to establish whether disease relevant geometries are indeed preferentially stabilized by the viral proteins. Further work is also needed to investigate the effect of FI10 and similar fragments on the intermediates such as dimers and other oligomers in the aggregation pathways that would seed fibril formation. This is especially important in the case of Aβ where such oligomers and not fibrils are the likely disease-causing agent.

## Supporting information

Atomic coordinates of conformations discussed in the article

## AUTHOR INFORMATION

## Author Contributions

**Mailnda B Premathilaka:** Formal analysis (equal); Investigation (equal); Visualization (lead); Writing – original draft (equal); **Ulrich H.E. Hansmann:** Conceptualization (lead); Funding acquisition (lead); Resources (lead); Supervision (lead); Writing – original draft (equal); Writing – review and editing (lead).

## Notes

The authors declare no competing financial interest.

## Data and Software Availability

The data supporting the results of this study are available in the Supporting Information and are publicly accessible at https://github.com/ouhansmannlab/amyloid_beta

## Supporting Information

**Supplementary Figures: SF1:** Ribbon representations of the three initial configurations of Aβ_1-42_ monomers and the two fibril models 7Q4B and 5KK3 bound with FI10 peptides in a 1:1 ratio; **SF2:** Average strandness as function of residues in the eight most populated clusters seen in the control simulations; **SF3:** Average strandness as function of residues in the 14 most populated clusters seen in the simulations where FI10 is present; **SF4:** Average number of intra-chain contacts between residues in the fibril conformations for the patient-derived fibril 7Q4B and the synthetic fibril 5KK3 as function of time. **Supplementary Tables: ST1:** Joint frequency (in percent) for pairs of the segments N1-C1, N1-C2, M1-C1 and M1-C2 being in a strand conformation; **ST2:** Frequency of Strandness for the segments N1, M1, C1 and C2 as function of the presence of the three salt bridges E1-H13, E22-K28 and D23-K28. **README** for a separate folder with atomic coordinates of selected Aβ_1-42_ monomers and fibril conformations discussed in the article.

## Acknowledgement

Our simulations were done using the SCHOONER cluster of the University of Oklahoma, ACCESS resources allocated under grant MCB160005 (National Science Foundation), and TACC resources allocated under grant under grant MCB20016 (National Science Foundation).

## Abbreviations

AD: Alzheimer’s disease
Aβ: Amyloid β
ɑS: ɑ synuclein
DSSP: Dictionary of secondary structure in proteins
HIV-TAT: Human immunodeficiency virus transactivator of transcription
HSV-1: Herpes simplex virus 1
MD: Molecular dynamics
PD: Parkinson’s disease
PDB: Protein data bank
PFC: Patient derived fibril-Control
PFF: Patient derived fibril with FI10
PME: Particle Mesh Ewald
RC1: Replica 1 Control
RC1_C1: Replica 1 Control-Cluster 1
RC2: Replica 2 Control
RC3: Replica 3 Control
RF1: Replica 1 with FI10
RF1_C1: Replica 1 with FI10-Cluster 1
RF2: Replica 2 with FI10
RF3: Replica 3 with FI10
Rg: Radius of gyration
RMSD: Root mean square deviation
RMSF: Root mean square fluctuation
SAA: Serum amyloid A
SARS-COV-2: Severe acute respiratory syndrome coronavirus-2
SASA: Solvent accessible surface area
SFC: Synthetic fibril control
SFF: Synthetic Fibril with FI10

