## Supplemental figures and tables for "Modulation of Aβ_1-42_ Aggregation by a SARS-COV-2 Protein Fragment"

#### Modulation of A$\beta_{1\text{-}42}$ Aggregation by a SARS-COV-2 Protein Fragment

Supporting information for this article consists of 2 tables and 4 figures listed on the following pages, and in a separate file (**AB-coordinates.zip**) a compressed folder with the atomic coordinates of the start and final configurations (as ASCII files in the PDB format) of all trajectories of the molecular dynamics simulations considered in this study. The folder contains also a README file to help identifying the various files.

### Table of content of supplementary information

**Figures:**

**SF1:** Ribbon representations of the three initial configurations of $A\beta_{1-42}$ monomers and the two fibril models 7Q4B and 5KK3 bound with FI10 peptides in a 1:1 ratio S3

**SF2:** Average strandness as function of residues in the eight most populated clusters seen in the control simulations. S4

**SF3:** Average strandness as function of residues in the 14 most populated clusters seen in the simulations where FI10 is present. S5

**SF4:** Average number of intra-chain contacts between residues in the fibril conformations for the patient-derived fibril 7Q4B and the synthetic fibril 5KK3 as function of time. S6

**Tables:**

**ST1:** Joint frequency (in percent) for pairs of the segments N1-C1, N1-C2, M1- C1 and M1- C2 being in a strand conformation. S7

**ST2:** Frequency of Strandness for the segments N1, M1, C1 and C2 as function of the presence of the three salt bridges E1-H13, E22-K28 and D23-K28. S8

**README for additional files in the compressed folder AB-coordinates.zip**

**(in separate file)** S9

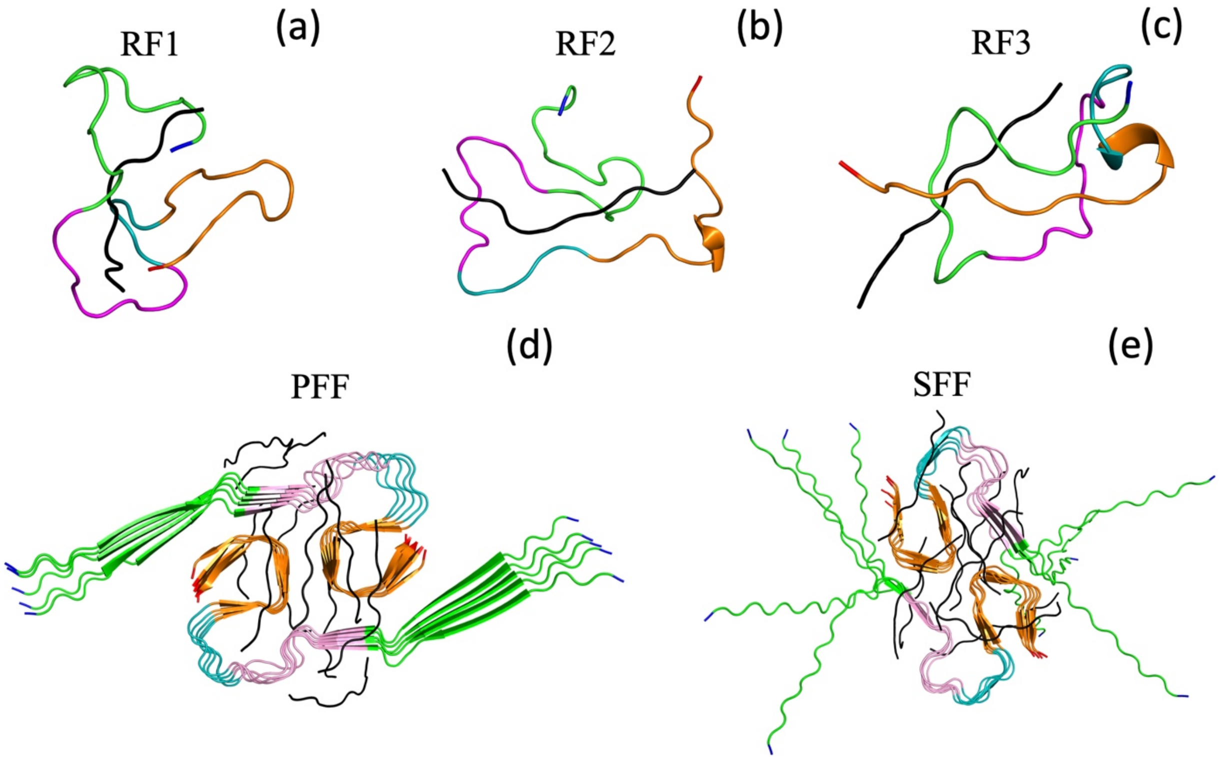


**Supplemental Figure SF1:** Ribbon representations of the three initial configurations (RF1, RF2, and RF3) of $A\beta_{1\text{-}42}$ monomers bound with a FI10 fragment (black) are shown in (a-c), with the various considered regions colored in green (region N, residues 1-15), pink (region M1, residues 16-24), teal (region M2, residues 25-29) and orange (region C, residues 30-42). The N- and C-terminals are colored in blue and red, respectively. Corresponding figures for the decamer $A\beta_{1\text{-}42}$ fibril models bound with ten FI10 chains used in our study are shown in (d) for the patient-derived model PFF with PDB-ID 7Q4B and in (e) for the synthetic fibril model SFF with PDB-ID 5KK3.

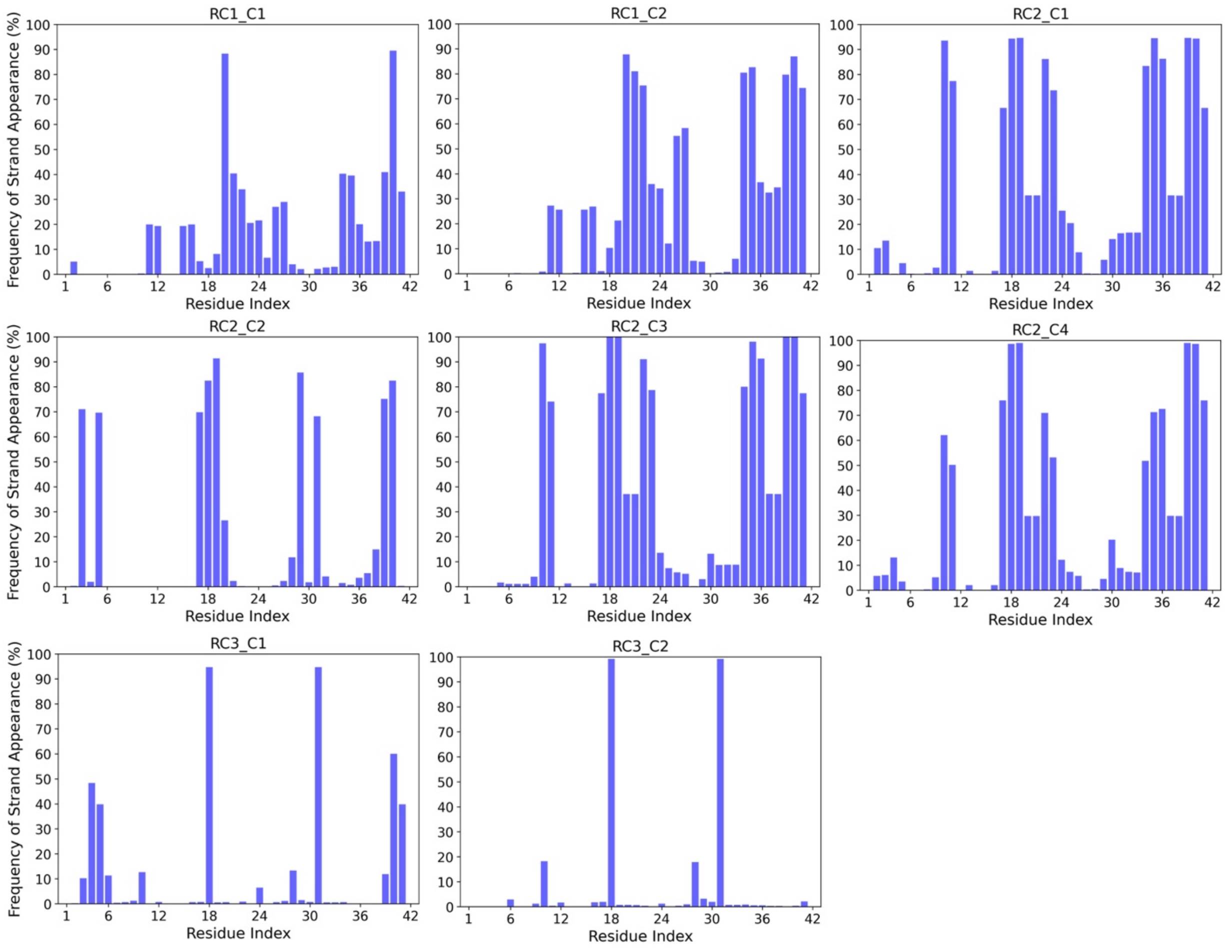


**Supplemental Figure SF2:** Average strandness as a function of residue number in the eight most populated clusters of configurations sampled over the final 2 μs in the three control trajectories (where FI10 is absent).

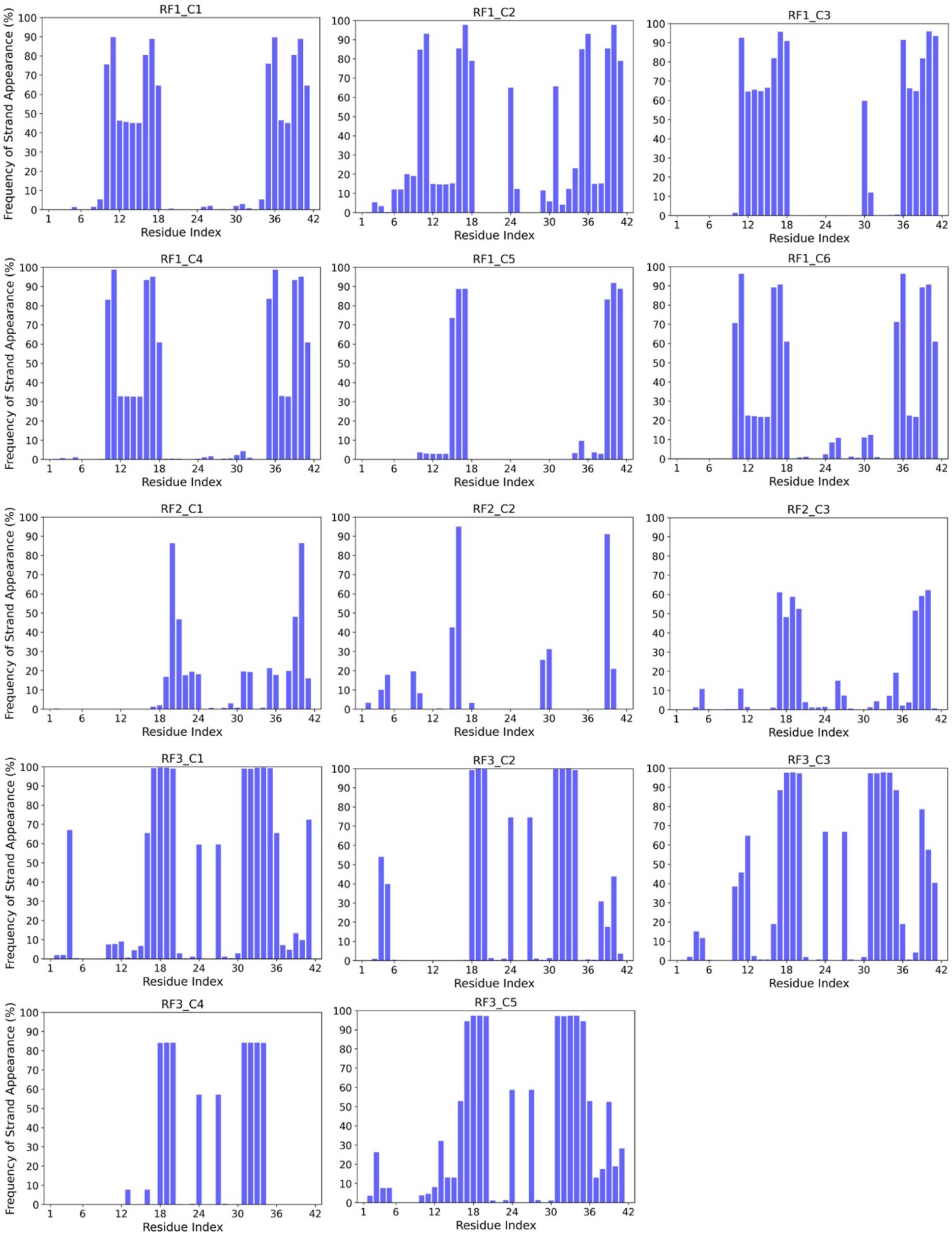


**Supplemental Figure SF3:** Average strandness as a function of residue number in the 14 most populated clusters of configurations sampled over the final 2 μs in the three trajectories where FI10 interacts with the A$\beta_{1\text{-}42}$ monomers.

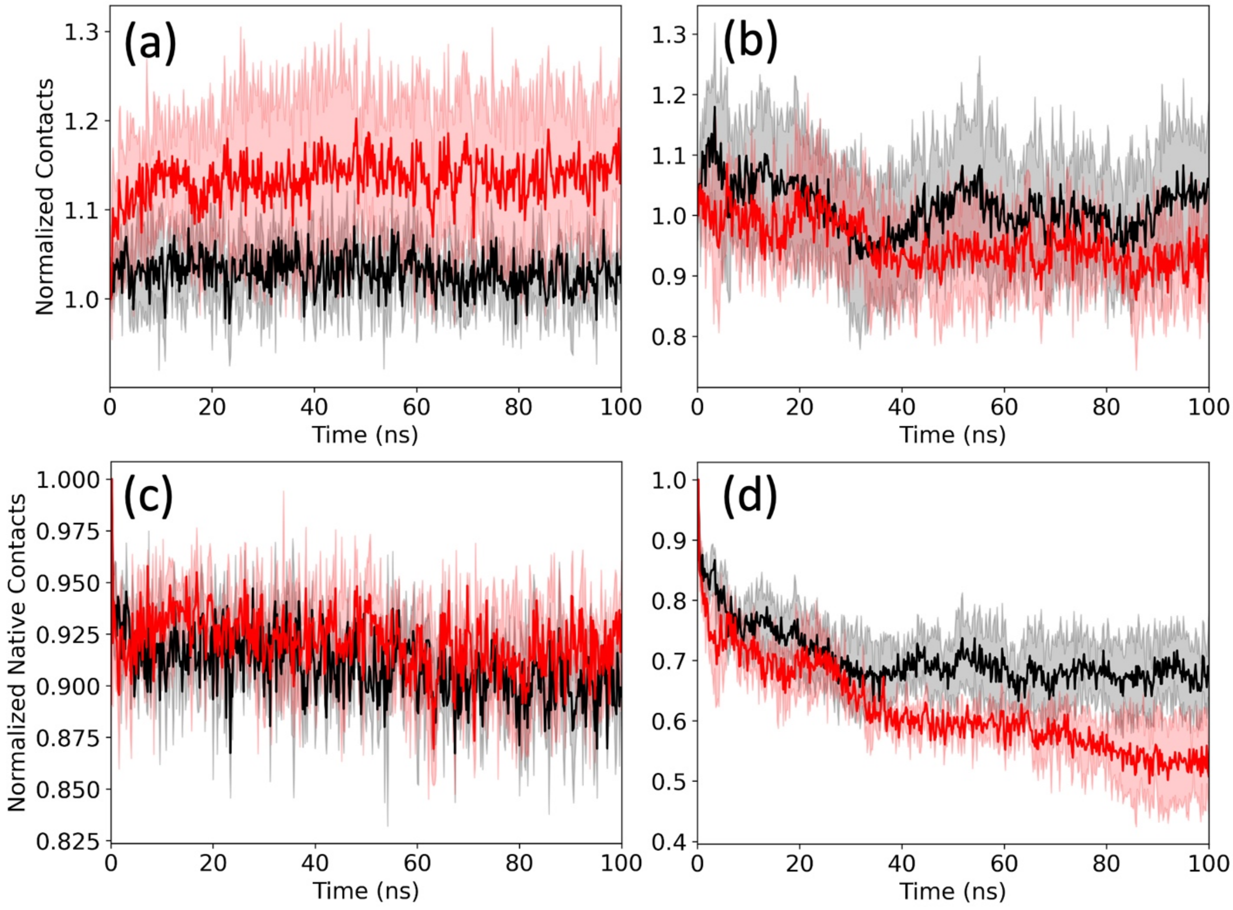


**Supplemental Figure SF4:** Average number of intra-chain contacts between residues in the fibril conformations for the patient-derived fibril 7Q4B (a) and the synthetic fibril 5KK3 (b) as function of time. Values for the control are drawn in black and the one from simulations with FI10 present in red. Averages are calculated over three independent trajectories and the corresponding standard deviations are given as shaded areas. For better comparison are the values normalized to unity for the conformation observed at 0.2 ns. In (c) and (d) we show the corresponding figures for native contacts, that is contacts which are seen in respective fibril models.

**Supplemental Table ST1** Joint frequency (in percent) for pairs of the segments N1-C1, N1-C2, M1- C1 and M1- C2 being in a strand conformation. Values are given for all considered clusters and averaged over all configurations sampled in the final two μs of each trajectory. Values are rounded to the closest integer. Standard deviation is given within the brackets.

| | Segments | | | |
|---|---|---|---|---|
| | N1∩C1 | N1∩C2 | M1∩C1 | M1∩C2 |
| Control | 0 (0) | 0 (0) | 25 (20) | 30 (30) |
| with FI10 | 4 (5) | 14 (16) | 34 (57) | 25 (24) |
| Clusters | | | | |
| RC1_C1 | 0 | 0 | 12 | 36 |
| RC1_C2 | 0 | 1 | 35 | 86 |
| RC2_C1 | 0 | 0 | 68 | 80 |
| RC2_C2 | 0 | 0 | 0 | 11 |
| RC2_C3 | 0 | 0 | 64 | 86 |
| RC2_C4 | 0 | 0 | 34 | 84 |
| RC3_C1 | 0 | 0 | 1 | 0 |
| RC3_C2 | 0 | 0 | 1 | 0 |
| RF1_C1 | 5 | 51 | 2 | 62 |
| RF1_C2 | 3 | 15 | 15 | 66 |
| RF1_C3 | 0 | 64 | 0 | 77 |
| RF1_C4 | 0 | 33 | 0 | 55 |
| RF1_C5 | 0 | 3 | 0 | 0 |
| RF1_C6 | 0 | 23 | 0 | 57 |
| RF2_C1 | 0 | 0 | 0 | 23 |
| RF2_C2 | 0 | 0 | 0 | 0 |
| RF2_C3 | 0 | 0 | 0 | 38 |
| RF3_C1 | 6 | 6 | 100 | 9 |
| RF3_C2 | 0 | 0 | 100 | 1 |
| RF3_C3 | 38 | 38 | 100 | 39 |
| RF3_C4 | 0 | 0 | 100 | 0 |
| RF3_C5 | 7 | 7 | 100 | 21 |

**Supplemental Table ST2:** Frequency of Strandness for the segments N1, M1, C1 and C2 as function of the presence of the three salt bridges E1-H13, E22-K28 and D23-K28.

| Frequency of Strand formation % | | | | | | | | |
|---|---|---|---|---|---|---|---|---|
| | Overall | | E11-H13 | | E22-K28 | | D23-K28 | |
| | Control | FI10 | Control | FI10 | Control | FI10 | Control | FI10 |
| N1 | 0.2(0.3) | 14(17) | 0 (0) | 27 (20) | 0.3 (0.3) | 5 (4) | 0.04 (0.05) | 12 (9) |
| M1 | 44(36) | 55(44) | 55 (22) | 57 (26) | 20 (23) | 27 (20) | 18 (14) | 87 (26) |
| C1 | 29(22) | 35(56) | 44 (16) | 19 (34) | 9 (18) | 7 (14) | 18 (13) | 82 (36) |
| C2 | 31(22) | 31(28) | 38 (23) | 47 (29) | 5 (1) | 38 (26) | 3 (9) | 17 (13) |
| Joint Probabilities % | | | | | | | | |
| N1∩C1 | 0 | 0 | 0 (0) | 2 (1) | 0.02 (1) | 1 (2) | 0 (0) | 8 (4) |
| N1∩C2 | 4(5) | 14(16) | 0 (0) | 26 (20) | 0.03 (2) | 5 (4) | 0.02 (2) | 11 (9) |
| M1∩C1 | 25(20) | 34(57) | 38 (17) | 17(35) | 9(18) | 6(14) | 16(12) | 82 (36) |
| M1∩C2 | 30(30) | 25(24) | 38(23) | 42(25) | 1 (2) | 25 (18) | 3(9) | 15 (11) |

**ReadMe** for additional files in the compressed folder **AB-coordinates (**available as separate file)

The folder **AB-coordinates** contains (besides this README.txt file) two folders (MONOMER and FIBRIL) that contain txt-files with the atomic coordinates (in PDB format) of AB(1-42) structures discussed in the article.

MONOMER;
This folder contains three sub-directories collecting AB(1-42) monomer configuration:

1) INITIAL:
Atomic coordinates (in PDB format) of the start-configurations of all six monomer simulations:
RC1_initial : Initial structure of AB(1-42) replica 1 control simulation
RC2_initial : Initial structure of AB(1-42) replica 2 control simulation
RC3_initial : Initial structure of AB(1-42) replica 3 control simulation
RF1_initial : Initial structure of AB(1-42) replica 1 with FI10 simulation
RF2_initial : Initial structure of AB(1-42) replica 2 with FI10 simulation
RF3_initial : Initial structure of AB(1-42) replica 3 with FI10 simulation

2) FINAL:
Atomic coordinates (in PDB format) of the final configurations of all six monomer simulations:
RC1_final : Final structure of AB(1-42) replica 1 control simulation
RC2_final : Final structure of AB(1-42) replica 2 control simulation
RC3_final : Final structure of AB(1-42) replica 3 control simulation
RF1_final : Final structure of AB(1-42) replica 1 with FI10 simulation
RF2_final : Final structure of AB(1-42) replica 2 with FI10 simulation
RF3_final : Final structure of AB(1-42) replica 3 with FI10 simulation

3) CENTROIDS:
This folder contains two sub-directories with cluster centroids

a) CONTROL:
Atomic coordinates (in PDB format) of the centroids of the clusters identified in the control simulations of the monomer, and discussed in the article:
RC1_C1 : Centroid structure of cluster 1 of replica 1 control simulation
RC1_C2 : Centroid structure of cluster 2 of replica 1 control simulation
RC2_C1 : Centroid structure of cluster 1 of replica 2 control simulation
RC2_C2 : Centroid structure of cluster 2 of replica 2 control simulation
RC2_C3 : Centroid structure of cluster 3 of replica 2 control simulation
RC2_C4 : Centroid structure of cluster 4 of replica 2 control simulation
RC3_C1 : Centroid structure of cluster 1 of replica 3 control simulation
RC3_C2 : Centroid structure of cluster 2 of replica 3 control simulation

b) FI10:
Atomic coordinates (in PDB format) of the centroids of the clusters identified in the simulations of the monomer interacting with FI10, and discussed in the article:
RF1_C1 : Centroid structure of cluster 1 of replica 1 with FI10 simulation
RF1_C2 : Centroid structure of cluster 2 of replica 1 with FI10 simulation
RF1_C3 : Centroid structure of cluster 3 of replica 1 with FI10 simulation
RF1_C4 : Centroid structure of cluster 4 of replica 1 with FI10 simulation
RF1_C5 : Centroid structure of cluster 5 of replica 1 with FI10 simulation
RF1_C6 : Centroid structure of cluster 6 of replica 1 with FI10 simulation

RF2_C1 : Centroid structure of cluster 1 of replica 2 with FI10 simulation
RF2_C2 : Centroid structure of cluster 2 of replica 2 with FI10 simulation
RF2_C3 : Centroid structure of cluster 3 of replica 2 with FI10 simulation
RF3_C1 : Centroid structure of cluster 1 of replica 3 with FI10 simulation
RF3_C2 : Centroid structure of cluster 2 of replica 3 with FI10 simulation
RF3_C3 : Centroid structure of cluster 3 of replica 3 with FI10 simulation
RF3_C4 : Centroid structure of cluster 4 of replica 3 with FI10 simulation
RF3_C5 : Centroid structure of cluster 5 of replica 3 with FI10 simulation

FIBRIL:
This folder contains two sub-directories collecting AB(1-42) fibril conformations:

1) 5KK3:
Atomic coordinates (in PDB-format) of the start and final configurations of the simulations of the synthetic fibril model with PDB-ID 5KK3:

a) CONTROL:
SFC1_initial : Initial structure of synthetic fibril control simulation replica 1
SFC1_final : Final structure of synthetic fibril control simulation replica 1
SFC2_initial : Initial structure of synthetic fibril control simulation replica 2
SFC2_final : Final structure of synthetic fibril control simulation replica 2
SFC3_initial : Initial structure of synthetic fibril control simulation replica 3
SFC3_final : Final structure of synthetic fibril control simulation replica 3

b) FI10:
SFF1_initial : Initial structure of synthetic fibril with FI10 replica 1
SFF1_final : Final structure of synthetic fibril with FI10 replica 1
SFF2_initial : Initial structure of synthetic fibril with FI10 replica 2
SFF2_final : Final structure of synthetic fibril with FI10 replica2
SFF3_initial : Initial structure of synthetic fibril with FI10 replica 3
SFF3_final : Final structure of synthetic fibril with FI10 replica 3

2) 7Q4B:
Atomic coordinates (in PDB-format) of the start and final configurations of the simulations of the patient-derived fibril model with PDB-ID 7Q4B:

a) CONTROL:
PFC1_initial : Initial structure of patient derived fibril control simulation replica 1
PFC1_final : Final structure of patient derived fibril control simulation replica 1
PFC2_initial : Initial structure of patient derived fibril control simulation replica 2
PFC2_final : Final structure of patient derived fibril control simulation replica 2
PFC3_initial : Initial structure of patient derived fibril control simulation replica 3
PFC3_final : Final structure of patient derived fibril control simulation replica 3

b) FI10:
PFF1_initial : Initial structure of patient derived fibril with FI10 replica 1
PFF1_final : Final structure of patient derived fibril with FI10 replica 1
PFF2_initial : Initial structure of patient derived fibril with FI10 replica2
PFF2_final : Final structure of patient derived fibril with FI10 replica 2
PFF3_initial : Initial structure of patient derived fibril with FI10 replica 3
PFF3_final : Final structure of patient derived fibril with FI10 replica 3
